# SARS-CoV-2 Nucleocapsid Protein forms biomolecular condensates and interacts with ALS-associated RNA binding proteins *in vitro*

**DOI:** 10.64898/2026.09.29.755449

**Authors:** Joseph-Patrick W.E. Clarke, Alexandra Keating, Veronica Noches, Crystal McLellan, Davide Dulcis, John Ravits, Michael J. Strong

**Affiliations:** Molecular Medicine Group, Robarts Research Institute, Schulich School of Medicine and Dentistry, Western University, London, ON, Canada; Department of Neurosciences, University of California San Diego School of Medicine, San Diego, CA, USA; Department of Clinical Neurological Sciences, Schulich School of Medicine and Dentistry, Western University, London, ON, Canada; Department of Pathology and Laboratory Medicine, Schulich School of Medicine and Dentistry, Western University, London, ON, Canada

**Keywords:** Keyword: Nucleocapsid Protein, Motor Neuron Disease, COVID-19, Neuronal Cytoplasmic Biomolecular Condensates, Optogenetics

## Abstract

As we have entered the endemic phase of COVID-19, there is a need to assess the long-term effects of SARS-CoV-2 infection on the trajectory of neurodegenerative disorders. Herein, we investigate whether the SARS-CoV-2 RNA-binding nucleocapsid protein (NCP; N-protein) may influence molecular processes relevant to motor neuron degeneration in amyotrophic lateral sclerosis (ALS). We detected N-protein immunoreactivity in ALS spinal cord motor neurons, including cytoplasmic N-protein-positive puncta that colocalized with cytoplasmic TAR DNA-binding protein 43 (TDP-43). In vitro, optogenetically induced N-protein biomolecular condensates (BMCs) showed reversible, liquid-like behavior and recovered after photobleaching. Using this model, we found that these condensates colocalized with TDP-43 and the RNA-binding proteins G3BP1/2, PABPC1 and TIA1. Condensate formation was associated with TDP-43 cytoplasmic mislocalization, altered *TARDBP* expression, and altered exon 3 splicing of *POLDIP3*. The N-terminal portion of N-protein influenced BMC formation, whereas truncated C-terminal proteoforms did not form detectable BMCs or show detectable TDP-43 colocalization in this assay. Extending these observations to a human cell model, we show that optogenetically induced N-protein BMCs colocalize with TDP-43 in human iPSC-derived motor neurons. Collectively, these findings support further investigation into potential molecular links between SARS-CoV-2 exposure and the trajectory of neurodegenerative disease.

## Introduction

Although the acute phase of the coronavirus disease 2019 (COVID-19) pandemic has passed, there is increasing evidence suggesting individuals exposed to the SARS-CoV-2 virus exhibit persistent neurological manifestations and may have a higher probability of developing neurodegenerative disease. For instance, current research has shown that neurological deficits worsened in Parkinson’s Disease patients during COVID-19, with new cases identified post COVID-19 at an increased rate (1), while in Alzheimer’s Disease, it has been demonstrated that COVID-19 is a significant risk associated with new onset (2). Additionally, while not associated with a specific neurodegenerative disease, studies have documented an increased risk of cognitive dysfunction, even two years after having contracted SARS-CoV-2, while other features of the post-COVID condition have resolved and are independent from the severity of the acute infection (3–7). To support these findings, there is evidence that patients who tested positive for SARS-CoV-2 show significant reductions in global brain size with cortical thinning (8). Overall, this postulate is further strengthened by previous observations on H1N1, as the SARS-CoV-2 pandemic is often compared to the 1918 H1N1 pandemic where Parkinsonian-like manifestations (i.e., encephalitis lethargica) were prominent (9, 10). Additionally, the peak age-distribution for SARS-CoV-2 infection was in the third through sixth decades, which constitutes a cohort of individuals most likely to harbour a latent neurodegenerative process that may be at risk for an accelerated neurodegenerative phenotype (2, 10, 11).

In the case of ALS, the evidence that SARS-CoV-2 infection can influence ALS onset or progression is, at best, indirect (2, 10). On the molecular level, while it is not entirely known how SARS-CoV-2 could modify such a process, there is a growing interest in the potential role of SARS-CoV-2 viral proteins participating in pathological biomolecular condensate (BMC) formation, which is a common neuropathological correlate for ALS and other neurodegenerative disorders (10, 12–14). ALS is the most common adult-onset neurodegenerative disorder of the CNS motor system, with many of the molecular manifestations observed in cortical and spinal cord motor neurons being attributed to the “loss-of-function” / “gain-of-function” aspects of the RNA-binding protein (RBP), TAR DNA-binding protein 43 (TDP-43), particularly in reference to changes in RNA metabolism (15, 16). Current evidence has shown that mislocalized cytoplasmic TDP-43 undergoes liquid-liquid phase separation (LLPS) into BMCs, often involving other RBPs, particularly with proteins involved in stress granule formation (17, 18).

Nucleocapsid protein (N-protein; NCP) is the most abundant structural SARS-CoV-2 protein expressed and acts as an RNA-binding protein involved in the transcription, translation and virion packaging of SARS-CoV-2 linear, positive-sense, single-stranded genomic RNA. Through an interaction with G3BP1, N-protein is involved in the process of stress granule formation (19). Like other RBPs, N-protein undergoes LLPS into BMCs and can form small homodimers and larger homo-oligomers (20–22). While the molecular consequences of N-protein BMCs are not fully described, current research suggests N-protein BMCs are important for RNA binding, expression on the cellular plasma membrane, extracellular secretion and induction of the host immune response (23, 24).

We have previously shown *in silico* a putative interaction between N-protein and TDP-43 that is reliant on the C-terminal portion of N-protein, thereby suggesting downstream altered TDP-43 molecular dynamics that may relate to the pathogenesis of ALS (25). To expand on this postulate, the current study examined N-protein immunoreactivity in human ALS spinal cord tissue and its molecular effects in cells *in vitro*. We report N-protein immunoreactivity in human ALS spinal cord motor neurons, including colocalization with mislocalized TDP-43. We further examine this association in immortalized cells (HEK293T) and human motor neurons [induced pluripotent stem cell (iPSC) derived] *in vitro* using an optogenetic N-protein condensation paradigm. In doing so, we demonstrate that the association profile of N-protein condensates extends beyond TDP-43, involving interactions with PABPC1, G3BP1, TIA1 and other proteins involved in splicing and translation. Downstream, we show that N-protein condensate formation is associated with TDP-43 cytoplasmic mislocalization and alterations in *TARDBP* expression and *POLDIP3* splicing, both TDP-43-regulated RNA-processing readouts. Finally, N-protein BMC formation and TDP-43 colocalization were not detected for previously described N-protein proteoforms (26).

Overall, this study examines SARS-CoV-2 N-protein in human postmortem ALS tissue and *in vitro* models and highlights the need for further investigation into whether SARS-CoV-2 exposure can alter the natural history of an existing neurodegenerative disease state and ultimately alter disease trajectories.

## Results

### N-Protein Immunoreactivity is Detected in Human ALS Spinal Cord Motor Neurons and Colocalizes with TDP-43

To expand our understanding of N-protein in biological systems and the effect SARS-CoV-2 infection has on neurodegenerative disease, we assessed N-protein immunoreactivity and its spatial relationship with TDP-43 in ALS spinal cord archival tissues. For samples obtained from the postmortem archive from the University Hospital, London Health Sciences Centre (London, Ontario, Canada) (N = 155), all samples were obtained using ethics research protocols approved by the Western University Health Sciences Research Ethics Board (HSREB) (HSREB #103735 & #124855). From the archive, we identified 21 cases with autopsy tissue obtained between 2009 and 2022 with a diagnosis of ALS [eight with pathological hexanucleotide expansion of C9*orf*72, three with SOD1 mutations (one each of A4T, I113T, D90A), and the remaining sporadic], including nine males and twelve females with a mean age of 62 ± 6.7 years and 65 ± 11.8 years (mean ± std. deviation), respectively (Supplemental Figure 1 A-C; Supplemental Table 1). We supplemented this with an additional eight sporadic ALS lumbar spinal cord samples (six males and two females with a mean age of 73 ± 8.6 years and 74 ± 3.5 years, respectively) obtained during the pandemic (provided by Target ALS CNS Repository Consortium; Dr. John Ravits, University of California San Diego, California, USA) (Supplemental Table 1) for a total of 29 ALS spinal cord samples. Six neuropathologically normal control cases (four males and two females with a mean age of 63 ± 6.0 years and both females aged 53) were also studied.

We initially tested the N-protein antibody used in this study on previously infected SARS-CoV-2 control lung tissue to test for positive N-protein reactivity, using both DAB and confocal IF methodologies (Supplemental Figure 1 D & E, respectively). In parallel, we also tested SARS-CoV-2 Spike, Membrane and Envelope immunoreactivity on infected control lung tissue (kind gift of Professor James Goldman, Columbia University, USA; data not shown). We confirmed that all nonSOD1 mutation ALS cases bore the neuropathological hallmark of the TDP-43 proteinopathy of ALS while the SOD1 cases demonstrated an upregulation of cytosolic TDP-43 with aggregate formation (Supplemental Figure 1 F & G).

To examine both N-protein and TDP-43 in ALS spinal cord tissue, we used confocal IF. We detected both N-protein and TDP-43 immunoreactivity in the cytoplasm of anterior horn spinal cord motor neurons from nine ALS patient spinal cord samples out of a combined total of 29 cases (Strong tissues N = 7 out of 21 cases; Ravits tissues N = 2 out of 8 cases). We did not detect nuclear N-protein in any of our samples (Figure 1 A-F). N-protein-positive cytoplasmic puncta had an average size of 0.85μm^2^ (Figure 1 G). Nuclear clearing of TDP-43 was observed in all ALS cases with isolated neuronal cytosolic inclusions of TDP-43 also observed in the majority (Figure 1 A-F). We, however, did observe mislocalized TDP-43 in all motor neurons (Figure 1 A-F). To quantify N-protein/TDP-43 colocalization, we calculated Pearson coefficients (PCC = 0.54) (Figure 1 B, D, F and H; Yellow box). We observed a positive, moderate colocalization that was further evident from 3D deconvolution imaging (Figure 1 B, D and F). Interestingly, the colocalization between N-protein and TDP-43 was not complete as we observed instances where cytoplasmic TDP-43 inclusions did not colocalize with N-protein (Figure 1 A, arrow with asterisk).

**Figure 1.**
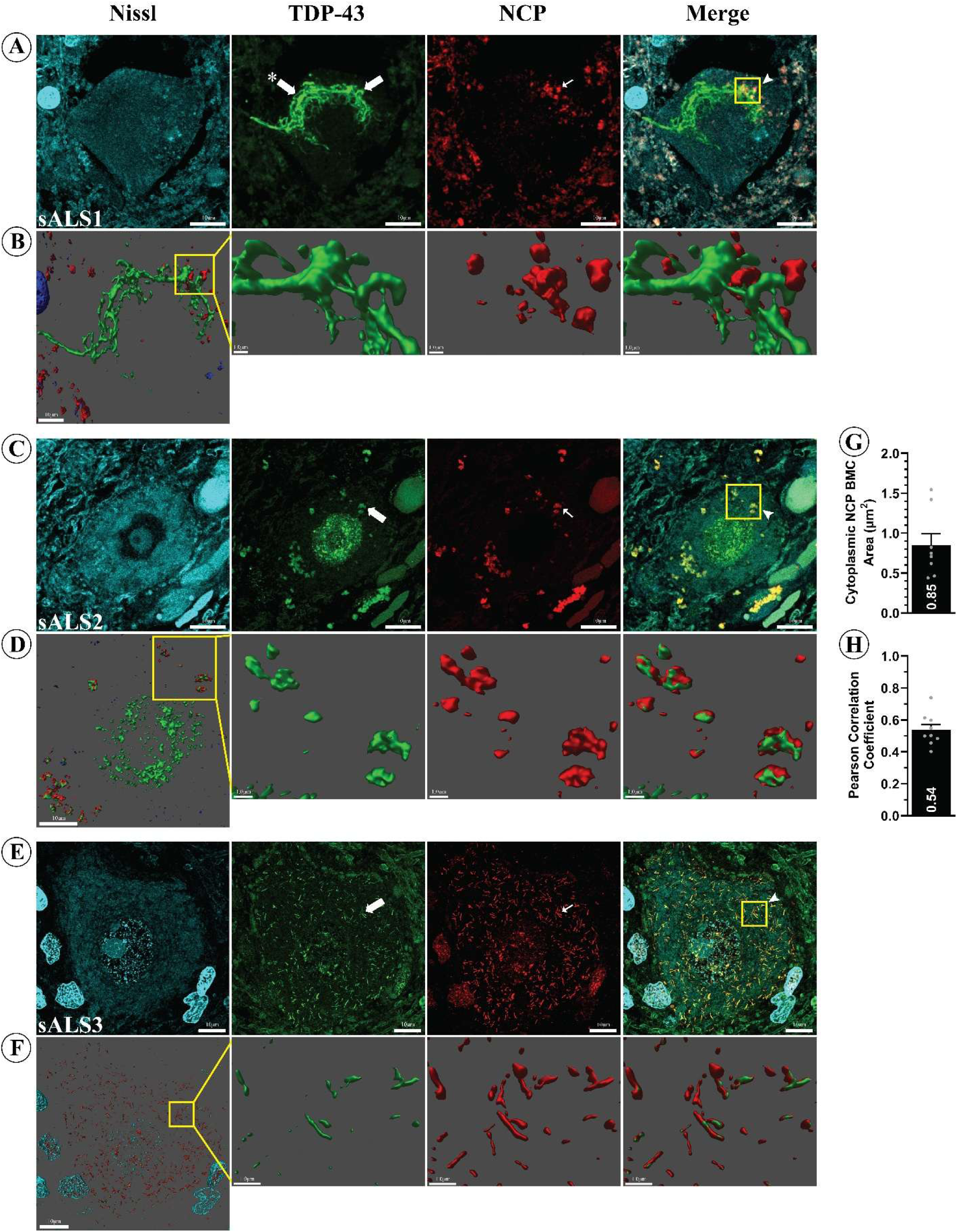
N protein is expressed in human ALS spinal cord motor neurons and interacts with cytoplasmic TDP-43. (A-F) Confocal IF from three of the nine positive N protein human patient ALS spinal cord cases indicating the presence of both increased cytoplasmic expression of TDP-43 and TDP-43 immunoreactive inclusions (Asterisk = cytoplasmic TDP-43 inclusion not colocalized with N protein). Also present is the expression of cytoplasmic N protein (Red fluorescence; small white arrows = cytoplasmic N protein), that shows colocalization with TDP-43 (Yellow box; arrowheads = colocalization between N protein and TDP-43). Nissl staining was performed to detect the nuclei of motor neurons (Blue fluorescence). Scale bars = 10.0μm. The colocalized N protein and TDP-43 in the merged images (yellow box) was 3D deconvoluted to remove background and further show interaction (B, D and F). Scale bars = 1.0μm. (G) The average size of cytoplasmic N protein BMCs measured 0.85 µm^2^. (H) Colocalization of N protein BMCs with TDP-43, using Pearson’s correlation coefficient analysis.

Cumulatively, these results indicate that a subset of ALS anterior horn motor neurons contains N-protein-positive cytoplasmic puncta that colocalize with cytoplasmic TDP-43.

### N-Protein Forms BMCs *in vitro*

To further explore the biological consequences of N-protein BMCs and their interaction with TDP-43 and other cytoplasmic proteins, we utilized an optogenetic system (Opto-NCP) (Supplemental Figure 2 A & B). For our live cell optogenetic paradigm, we exposed transiently transfected HEK293T cells with 465 nm BL for 60 minutes to induce Opto-NCP self-association, with quantification of cytoplasmic Opto-NCP BMCs every 10 minutes during stimulation (Figure 2 A & B). We demonstrated that Opto-NCP BMC formation is specifically facilitated by BL stimulation of the Opto-tagged protein by comparing the self-association dynamics of Opto-NCP with mNCP-aCh (control construct lacking the Opto tag) with the latter failing to form condensates (Figure 2 A & B). To confirm that the observed effect of BL stimulation on Opto-NCP BMC biophysics is defined by the properties of N-protein, and not due to the Opto or mCherry tags, we compared the effect of BL stimulation between Opto-NCP and mCh-Opto (lacking the N-protein gene) (Figure 2 A & B). We found that Opto-NCP association dynamics are distinct from mCh-Opto, where Opto-NCP formed cytoplasmic BMCs at a half-maximal kinetic association rate (KA_1/2MAX_) of 16.6 minutes, compared to a KA_1/2MAX_ of two minutes for mCh-Opto (Figure 2 B).

**Figure 2.**
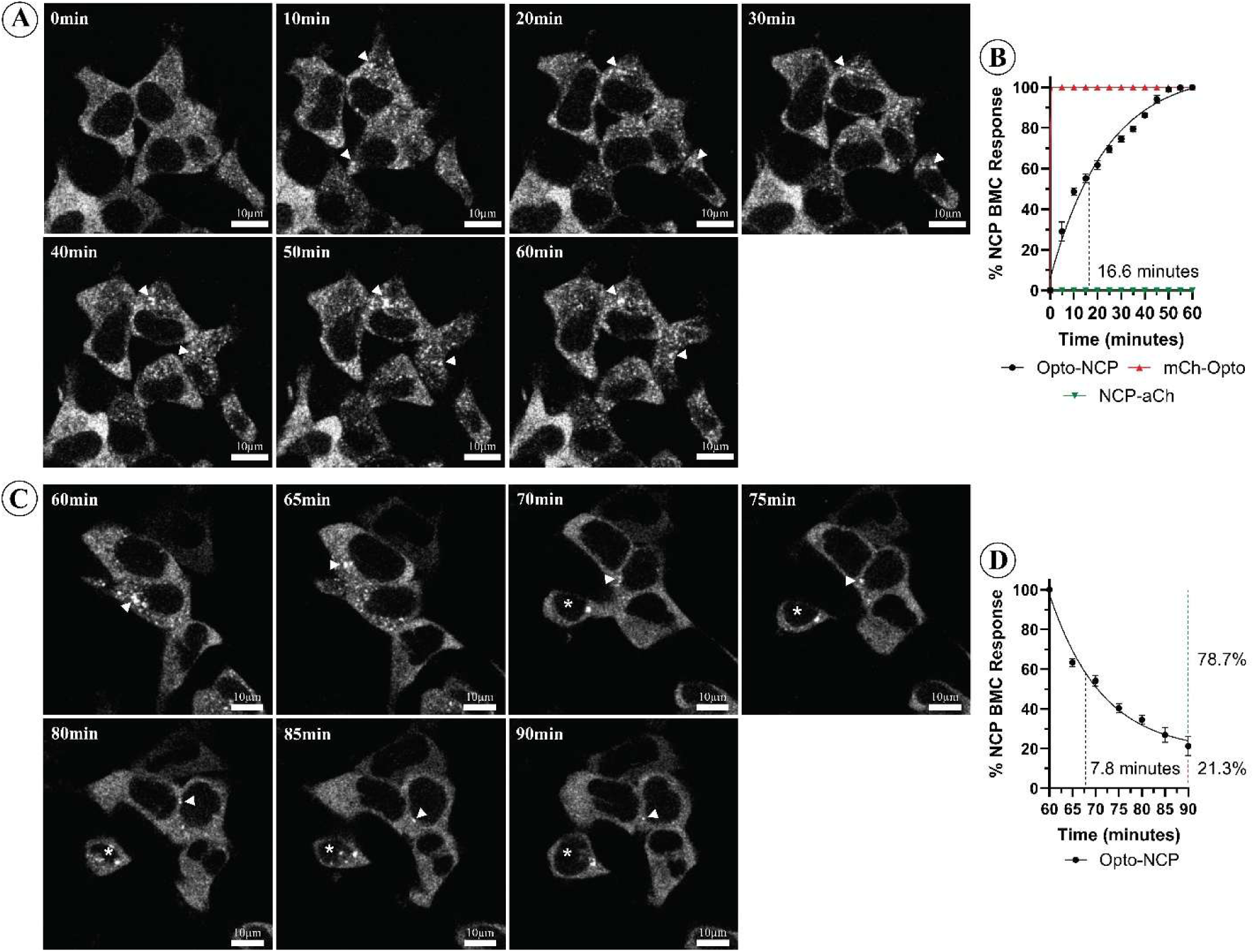
*In vitro* Opto-NCP undergoes LLPS with BL stimulation. (A) Representative images of Opto-NCP BMC formation during a live cell BL stimulation protocol. HEK293T cells expressing Opto-NCP were exposed to BL stimulation (10,000 lux, 465 nm) for 60 minutes, with images acquired every 10 minutes. Arrowheads indicate the formation of Opto-NCP BMCs. Scale bars = 10μm. (B) Time course quantification of BMC formation during 60-minute BL stimulation for each construct (Opto-NCP, mCh-Opto and NCP-aCh), resulting in a kinetics curve for association dynamics. Dashed line indicates KA_1/2Max_. Data shown are mean ± S.E.M. for three biological replicates. (C) Representative images Opto-NCP BMC dissolution after a live cell BL stimulation protocol. After 60 minutes of BL stimulation, the BL stimulation was suspended for 30 minutes, with images acquired every 10 minutes. Arrowheads indicate Opto-NCP BMCs. Asterisk indicates a cell with BMCs that did not dissociate after BL stimulation suspension. Scale bars = 10.0μm. (D) A time-course measurement of the rate of BMC disappearance following cessation of BL stimulation yielded a KD_1/2Max_ of 7.8 minutes (black dashed line). 30 minutes following cessation of BL stimulation, 78.7% of BMCs resolved (Green dashed line) while 21.3% of the initial BMCs remained (Red dashed lined), suggesting a more stable fraction resistant to further dissolution within the experimental time window. Data shown are mean ± S.E.M. for three biological replicates.

In our paradigm, we also analyzed the dissociation dynamics of Opto-NCP BMCs for 30 minutes after BL stimulation (90 minutes total experimentation time), again quantifying cytoplasmic Opto-NCP BMCs every 10 minutes (Figure 2 C & D). This analysis allows for the interpretation of BMC dissolution and overall solubility. From this paradigm, we found that Opto-NCP BMCs readily dissociated after BL stimulation, with a half-maximal kinetic dissociation rate (KD_1/2MAX_) of 7.8 minutes (Figure 2 C and D). Of note, 78.7% ± 4.8% (S.E.M.; 95% CI) of all cells underwent total BMC dissociation, while 21.4% ± 4.8% (S.E.M.; 95% CI) of all cells still contained BMCs at the end of experimentation (Figure 2 D; Asterisk = cell expressing Opto-NCP BMCs) suggesting a more stable pool of N-protein oligomers.

To further directly assess the solubility and molecular dynamics of Opto-NCP BMCs, we performed fluorescence recovery after photobleaching (FRAP) on individual BMCs (Figure 3 A). Quantitative analysis showed a 48.9% recovery of fluorescence intensity after FRAP, indicating that the BMCs possessed liquid-like characteristics (Figure 3 B). From our analysis, we also deduced that the BMCs, while liquid-like, are structured, as the diffusion coefficient was slow (0.029 μm^2^/second) (Figure 3 B). Consistent with our earlier observations, this structured characteristic is further emphasized by the similarity of the mobile and immobile fractions (44.3% vs. 55.8%, respectively), with the mobile fraction containing freely diffusible molecules, and the immobile fraction containing molecules that remain spatially confined (Figure 3 B).

**Figure 3.**
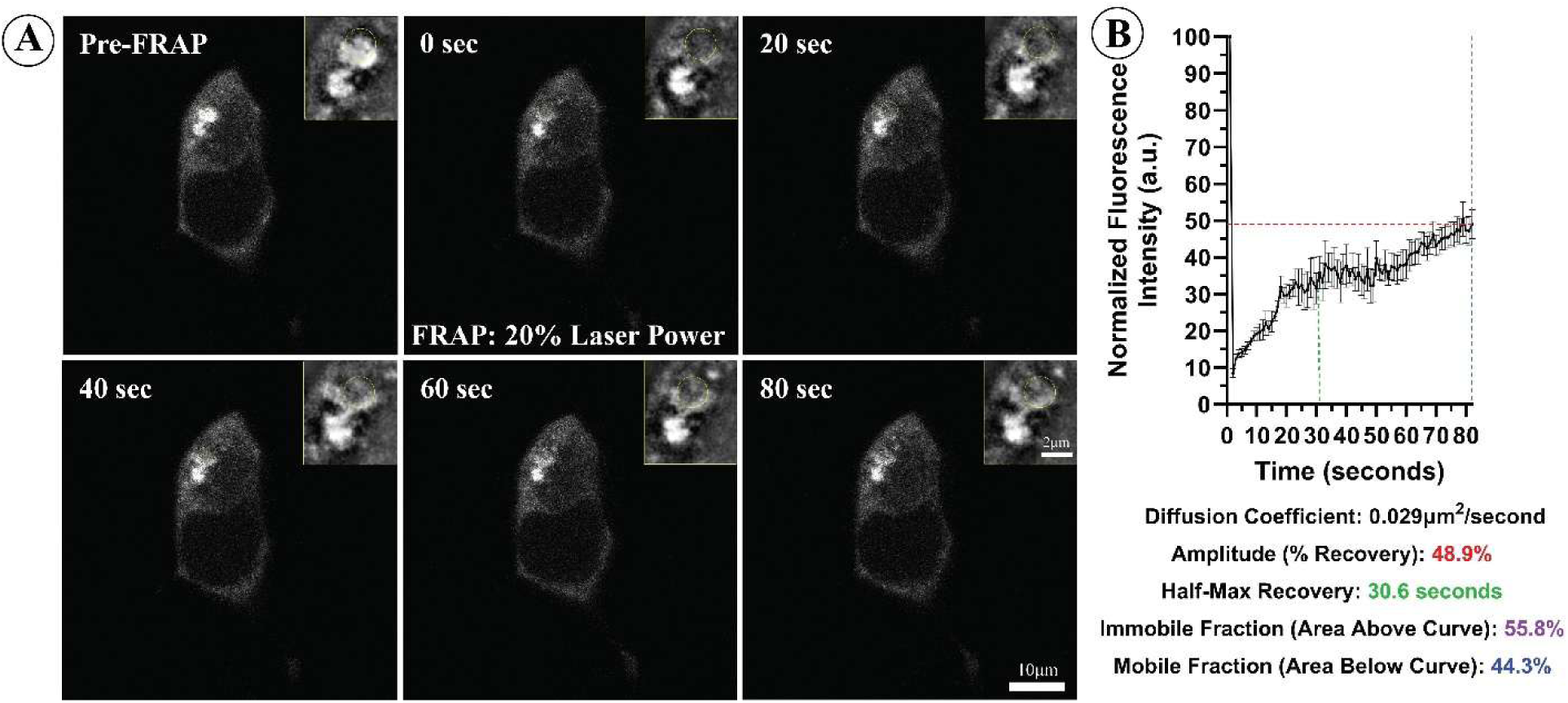
BL-stimulated *in vitro* Opto-NCP BMCs are soluble. (A and B) Representative images and quantification of Opto-NCP BMC FRAP. After 60 minutes of BL stimulation to form Opto-NCP BMCs, FRAP (20% laser bleach for 1.7 seconds in a circular 2.0µm^2^ area partially encompassing an Opto-NCP BMC) was performed on individual BMCs. Yellow boxes = magnification of FRAP areas. Yellow circles = 2.0µm^2^ area of FRAP. Data shown are mean ± S.E.M. for three biological replicates. Scale bars = 10.0μm and 2.0μm.

Overall, these observations show that the Opto-NCP construct forms reversible condensates with liquid-like properties under blue-light stimulation, providing a controllable model of N-protein condensation consistent with other optogenetic condensate systems (27–30).

### BMCs of N-Protein Colocalize with ALS-Associated RNA Binding Proteins *in vitro*

Having observed the colocalization of N-protein and TDP-43 in a subset of ALS patient spinal cord samples, we next investigated whether this colocalization could be recapitulated *in vitro* and whether it extended to other components of the ribonucleotide protein (RNP) complex. Following blue light stimulation of Opto-NCP as described above in HEK293T cells, using confocal microscopy we observed colocalization between N-protein and TDP-43 (PCC = 0.31) or pathologically phosphorylated (phospho-TDP; PCC = 0.31) (Figure 4 A - D). In control experiments, we observed no colocalization of TDP-43 in the presence of the empty mCh-Opto or non-Opto N-protein plasmids (Supplemental Figures 3 & 5).

**Figure 4.**
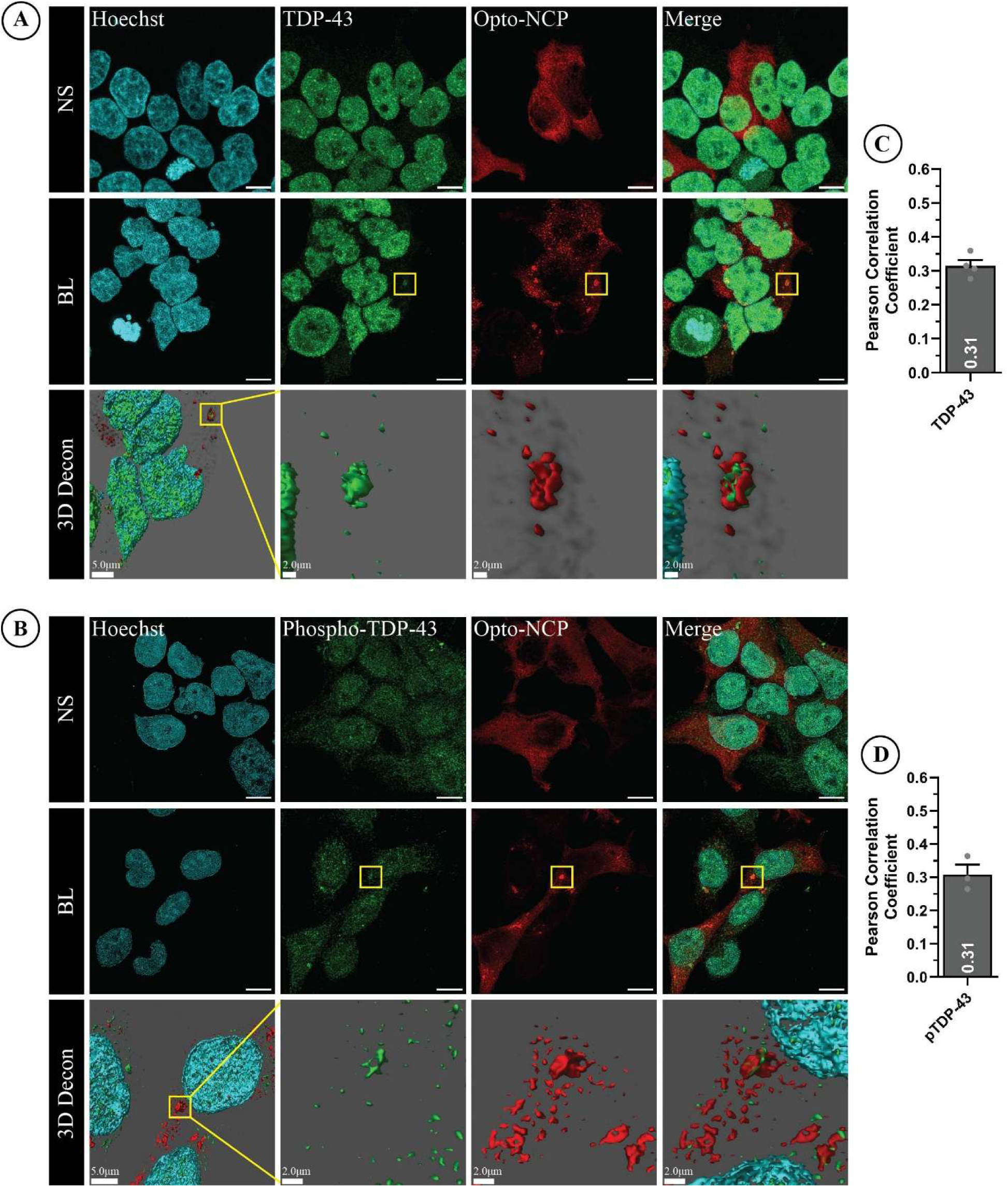
*In vitro* Opto-NCP BMCs colocalize with TDP-43 and phospho-TDP-43. BL induced Opto-NCP BMCs colocalize with TDP-43 (PCC = 0.31) (A and C) and phosphorylated-TDP-43 (PCC = 0.31) (B and D). BL = Blue light stimulation. NS = No stimulation. The Opto-NCP BMCs had an average size of 3.19µm^2^ (E). Yellow box = formation of NCP BMCs and colocalization with TDP-43. Scale bars = 10.0μm. Merged images underwent 3D deconvolution to remove background and improve the visual specificity of colocalization. Scale bars = 5.0μm and 2.0μm.

To determine whether the association of TDP-43 with NCP BMCs reflected participation in a larger RNP complex, we analyzed a subset of RNA binding proteins associated with ALS for colocalization with Opto-NCP BMCs using our optogenetic paradigm. Using confocal microscopy, we observed specific colocalization with PABPC1 (PCC = 0.47) (Supplemental Figure 4 A and B), G3BP1 (PCC = 0.43) (Supplemental Figure 4 C and D) and TIA1 (PCC = 0.43) (Supplemental Figure 4 E and F). In all cases, we 3D deconvoluted the results to remove background and improve the visual specificity of colocalization. Colocalization by confocal imaging was not observed for either FUS or hnRNP A1 (data not shown).

To determine whether our results were influenced by any cell stress that may be caused by our BL stimulation paradigm, we examined cell stress using two independent visualization methodologies. First, we examined the colocalization of eIF2S1 and phosphorylated eIF2S1 (phospho-eIF2S1) to G3BP1-positive N-protein BMCs (Supplemental Figure 5 A and B) [interaction previously described in the literature (31, 32)]. We analyzed cells that were a) transfected with Opto-NCP, b) formed Opto-NCP BMCs with BL stimulation and c) were positive for G3BP1 colocalization with N-protein BMCs. When we analyzed both eIF2S1 and phospho-eIF2S1, we found no colocalization of either with N-protein or G3BP1, suggesting that these markers did not detect a stress response under our conditions (Supplemental Figure 5 A and B). Additionally, we performed a Comet assay that sensitively measures DNA damage in individual cells and can be used to assess DNA damage-related cellular stress (Supplemental Figure 5 C, D and E). In this experimentation, we expanded our BL stimulation to six hours to chronically saturate our samples with BL. Even within this expanded timeframe, our data show minimal DNA tail formation (i.e., DNA tail length and tail DNA percentage) in the BL stimulated samples that was not significantly different from the paired non-stimulated samples (Supplemental Figure 5 C, D and E). Positive control samples included cells treated with 0.4 mM methylglyoxal for 24 hours in which robust DNA tail formations were observed with Comet assay (data not shown). Together, these results indicate that BL stimulation did not measurably increase eIF2S1/phospho-eIF2S1 recruitment or DNA damage under the conditions tested.

To further expand on our N-protein interaction network beyond a few measurable targets, we next performed a proteomic analysis on co-immunoprecipitated samples targeting the Opto-NCP with an mCherry antibody from non-stimulated and BL stimulated cell lysates followed by mass spectrometry of the immunoprecipitate (Figure 5 A). From this analysis, we identified increased association of TDP-43 and G3BP1 with N-protein following BL stimulation (Figure 5 A). We also identified association with G3BP2, components of both the 40S (RSP19 and RPS26) and 60S (RPL5 and RPL6) ribosomal subunits, as well as the Serine/Arginine-rich splicing factor 1 (SRSF1), DNA polymerase delta interacting protein 3 (POLDIP3) and, in contrast to confocal microscopy, FUS (Figure 5 B). To a lesser extent, we also identified associations with PABPC1, hnRNP A1, hnRNP A3, hnRNP H2, eukaryotic translation elongation factor 1 epsilon 1 (EEF1E1) and eukaryotic translation initiation factor subunit 3 (eIF3C) (Figure 5 B).

**Figure 5.**
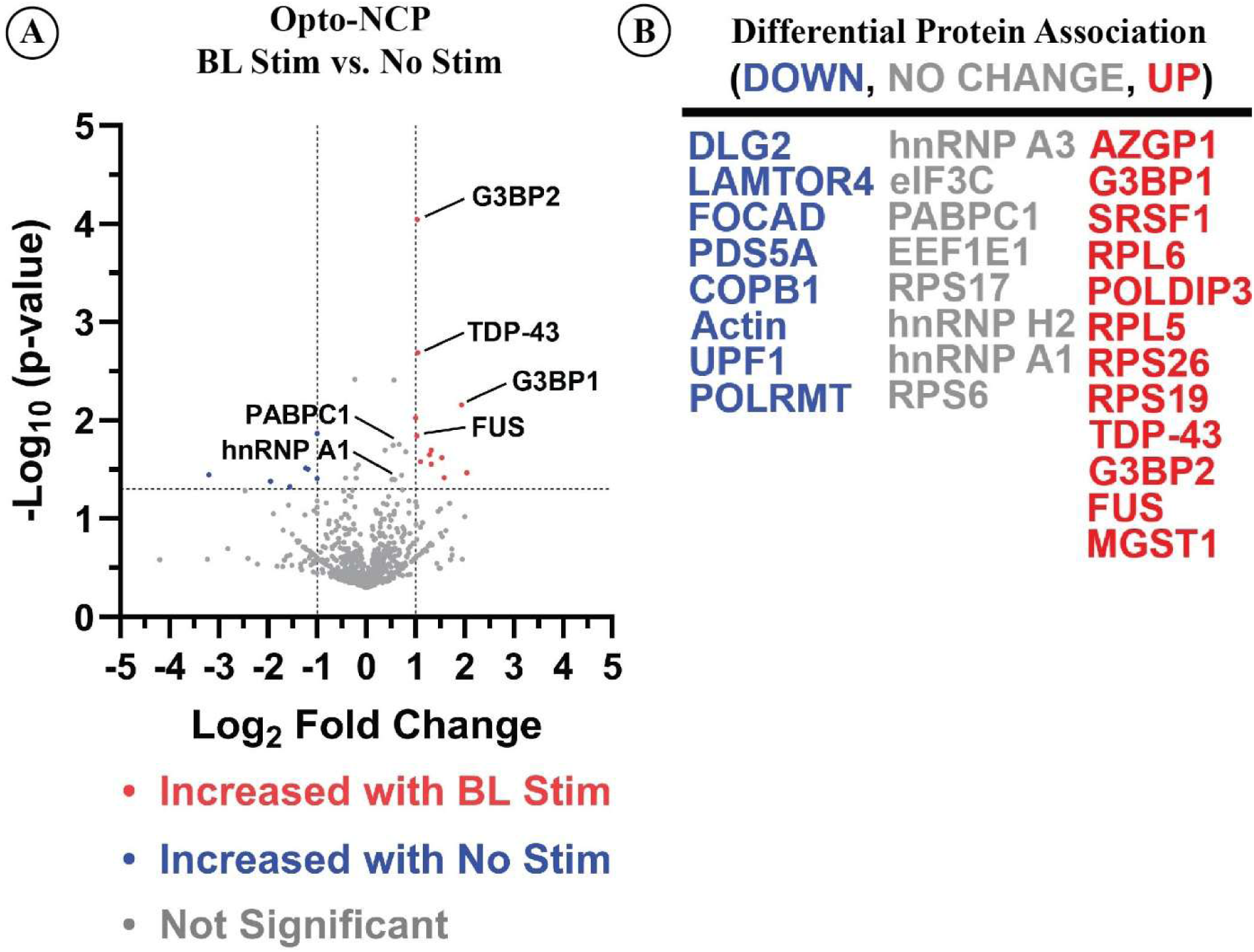
Differential protein expression associated with Opto-NCP following BL stimulation. Following BL stimulation in the presence of Opto-NCP, immunoprecipitation of the NCP BMCs was performed using the NCP mouse monoclonal antibody followed by mass spectroscopy of the immunoprecipitant. (A and B) Mass spectrometry confirmed the interactions between N protein with TDP-43, PABPC1 and G3BP1, and outlines other interactions with proteins involved in transcription, splicing and translation.

Overall, these results support association of N-protein with TDP-43 and expand the candidate N-protein-associated network to other RNPs, particularly proteins involved in transcription, splicing and translation.

### N-Protein BMCs Affect TDP-43 Mislocalization *in vitro* and Are Accompanied by RNA Metabolic Changes

From our confocal microscopy experimentation, we observed a significant nucleocytosic shift in the subcellular localization of TDP-43 with a reduction in nuclear TDP-43 (Figure 6 A and B). To determine whether the mislocalization of TDP-43 associated with N-protein condensate formation is accompanied by molecular effects, we next measured changes in RNA metabolism through the expression of TDP-43 RNA targets. Specifically, using expression RT-PCR, we measured the RNA changes in *TARDBP* and *POLDIP3*. TDP-43 is highly regulated and autoregulates *TARDBP* mRNA through alternative splicing events concentrated within exon 2, exon 6, and the 3′ untranslated region (3′UTR) (33). This autoregulation is known to be perturbed in ALS, and thus *TARDBP* is a specific marker RNA to assess TDP-43 function (33). For *POLDIP3*, TDP-43 has been shown to promote the inclusion of Exon 3 during RNA splicing, producing a functional, longer *POLDIP3* mRNA transcript (34, 35). Our results show a significant reduction in *TARDBP* expression (Figure 6 C and D) and a significant increase in the expression of a shorter *POLDIP3* RNA transcript, indicating the loss of Exon 3 during RNA splicing (Figure 6 C and E).

**Figure 6.**
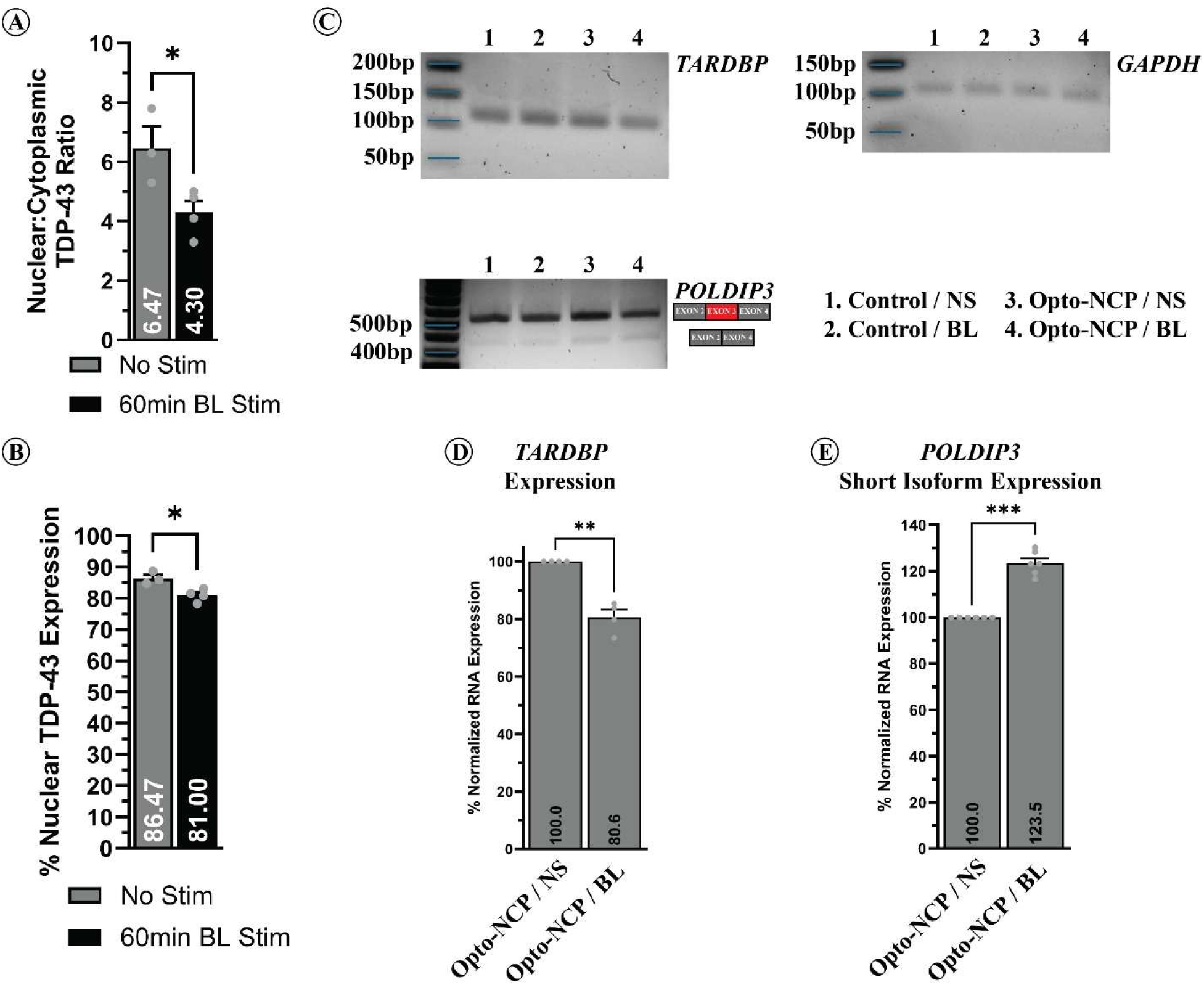
TDP-43 interaction with Opto-NCP causes its mislocalization and perturbs RNA metabolism in HEK293T cells. (A) Co-immunoprecipitation (Co-IP) of non-stimulated (Lanes 1, 3, 5) and BL stimulated (Lanes 2, 4, 6) Opto-NCP samples demonstrates interaction between N protein and TDP-43. (B) Densitometric analysis of Co-IPs in (A). (C and D) Following BL stimulation, mislocalization of TDP-43 in BL was measured by the nuclear:cytoplasmic ratio of TDP-43 and the percent nuclear amount of TDP-43 was observed. Assessing the RNA expression of *TARDBP* and the cryptic exon expression of *POLDIP3* in cells expressing Opto-NCP BMCs (non-stimulated and BL stimulated), a significant decrease in total *TARDBP* RNA (E and F) and a significant increase in the expression of a shorter *POLDIP3* cryptic exon (E and G) in BL stimulated Opto-NCP cells was observed. \**p* < 0.05; \*\**p* < 0.01; \*\*\**p* < 0.001; 95 % Confidence Interval.

Cumulatively, these results show the N-protein/TDP-43 interaction is accompanied by changes in TDP-43 localization and RNA-processing readouts consistent with altered TDP-43 function.

### N-Protein Proteoforms Attenuate BMC Formation and TDP-43 Colocalization

Our current results describe an association between full-length N-protein and TDP-43. However, N-protein can exist as shorter C-terminal proteoforms, NCP^M*210^ and NCP^M*234^ (26). To address this, we expressed the N-protein proteoforms (Opto-NCP_M210_ and Opto-NCP_M234_) using the optogenetic system in HEK293T cells and determined their colocalization with TDP-43 following BL stimulation (Figure 7 A). Neither N-protein proteoform formed detectable BMCs (Figure 7 B and C). This suggests that the N-terminal portion of N-protein (amino acid 1-209) may play a role in both processes. To ascertain the effect of the N-terminal portion of N-protein on its ability to form BMCs and colocalize with TDP-43, we optogenetically tested the N-terminal portion of N-protein in isolation (Opto-NCP_1-209_) (Figure 7 A). Like full-length N-protein, the NCP_1-209_ formed BMCs. However, they did not colocalize with TDP-43 (Figure 7 D; Supplemental Figure 3). This is consistent with our prior work showing that the interaction between N-protein and TDP-43 requires the C terminus domain of N-protein (25).

**Figure 7.**
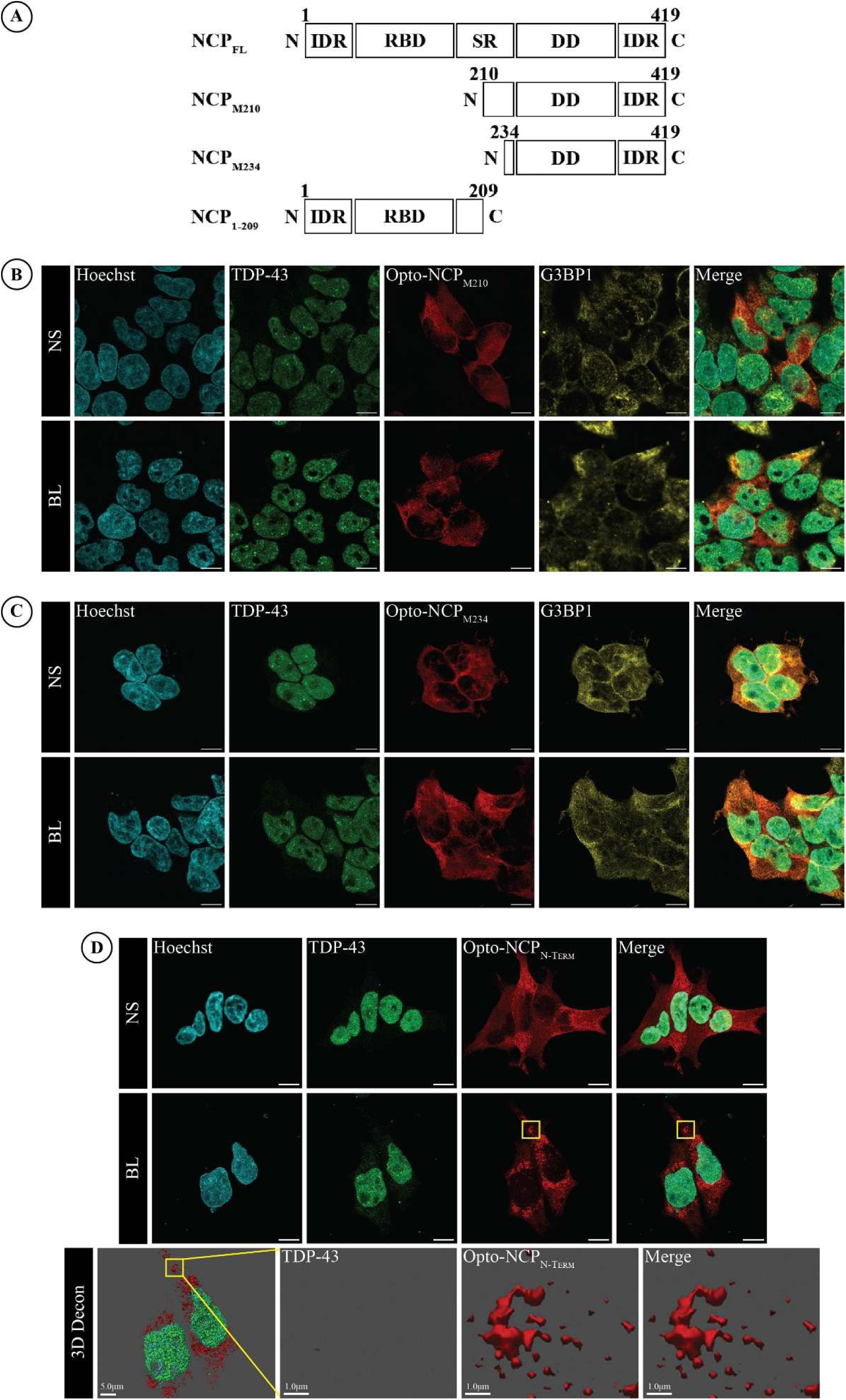
The N-terminal portion of N protein influences its ability to form BMCs. (A) Illustration of the full-length (NCP_FL_), NCP_M210_ and NCP_M234_ proteoforms and truncation of NCP (NCP_1-209_) used in this study. Amino acid positions are labelled relative to full-length NCP. IDR = intrinsically disordered region. RBD = RNA binding domain. SR = serine-arginine-rich domain. DD = dimerization domain. (B and C) No BMCs were formed following BL stimulation in the presence of either Opto-NCP_M210_ and Opto-NCP_M234_. Scale bars = 10.0μm. (D) Expressing Opto-NCP_1-209_ in contrast resulted in BMC formation following BL stimulation. Of note however, Opto-NCP_1-209_ BMCs did not colocalize with TDP-43. BL = Blue light stimulation. NS = No stimulation. Yellow box = formation of N protein BMCs and colocalization. Scale bars = 10μm. Merged images underwent 3D deconvolution to remove background and improve the visual specificity of colocalization. Scale bars = 5.0μm and 1.0μm.

Overall, these results suggest that distinct N- and C-terminal regions contribute to N-protein BMC formation and TDP-43 colocalization in this optogenetic assay.

### N-Protein BMCs Colocalize with TDP-43 in Cultured iPSC Motor Neurons

We next performed our optogenetic experimental paradigm in mature human induced pluripotent stem cell (iPSC) motor neurons using lentiviral transduction *in vitro*. We first created third generation lentiviral particles containing Opto-NCP, and transduced iPSC motor neurons, followed by a five-day incubation allowing for the expression of Opto-NCP. Our results show expression of optogenetically induced N-protein biomolecular condensates in iPSC motor neurons, and with BL stimulation, the formation of N-protein BMCs (Figure 8 A). Like our previous *in vitro* cell culture results, we observed colocalization of TDP-43 (PCC = 0.37) (Figure 8 A and B) and phospho-TDP-43 (PCC = 0.37) (Figure 8 C and D) with N-protein BMCs in the iPSC motor neuron cultures.

**Figure 8.**
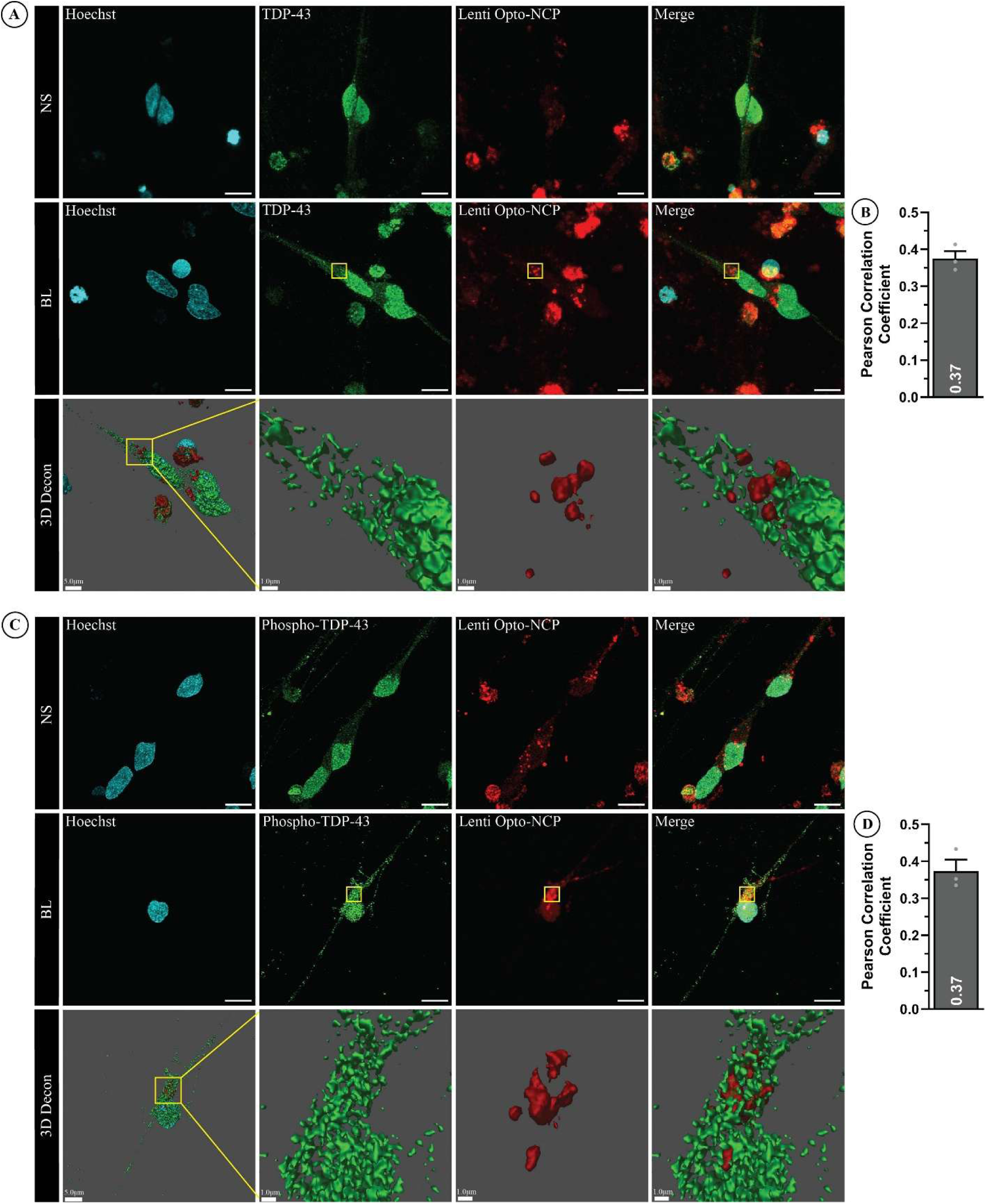
Opto-NCP BMCs colocalize with TDP-43 and phospho-TDP-43 in *in vitro* human iPSC motor neurons. Cultured iPSC motor neurons were matured for seven days *in vitro* before transduction with Opto-NCP lentivirus for five days. Blue light induced Opto-NCP BMCs form in human motor neurons and colocalize with both TDP-43 (PCC = 0.37) (Figure 8 A and B) and phosphorylated-TDP-43 (PCC = 0.37) (Figure 8 C and D). BL = Blue light stimulation. NS = No stimulation. Yellow box = formation of NCP BMCs and colocalization. Scale bars = 10.0μm. Merged images underwent 3D deconvolution to remove background and improve the visual specificity of colocalization. Scale bars = 5.0μm and 1.0μm.

Collectively, these results show that Opto-NCP forms BMCs that colocalize with TDP-43 in human iPSC-derived motor neurons, establishing a human motor-neuron model for future mechanistic studies.

## Discussion

In this study, we detected N-protein immunoreactivity in cytoplasmic puncta that colocalized with TDP-43 in spinal motor neurons from human ALS postmortem tissue spanning pre-pandemic and pandemic periods. We further show in cells *in vitro* that optogenetically induced N-protein BMCs are liquid-like and display aspects of LLPS, having both biophysical dynamics of association and dissociation and diffusion of particles within condensates. We further describe the colocalization of N-protein with TDP-43, phospho-TDP-43, PABPC1, G3BP1 and TIA1, as well as proteins involved in transcription, splicing and translation. Expanding on its association with TDP-43, we found that N-protein condensate induction was accompanied by TDP-43 cytoplasmic mislocalization, decreased *TARDBP* expression, and altered *POLDIP3* exon 3 splicing. In parallel, we also examined the *in vitro* influence of N-protein proteoforms (i.e., NCP_M210_ and NCP_M234_) on their ability to form BMCs, and whether these affect TDP-43 colocalization. We demonstrate that the N-protein proteoforms do not form BMCs, and do not colocalize with TDP-43, while the N-protein N-terminus (amino acids 1-209) alone forms coarse granular BMCs, but with absent TDP-43 colocalization. These results suggest that both N-protein oligomerization and C-terminal determinants contribute to TDP-43 colocalization in this system. Finally, to recapitulate and expand on our findings in a human motor-neuron model, we utilized human induced pluripotent stem cell motor neurons transduced with lentiviral optogenetic N-protein. Using this model, we show that N-protein BMCs colocalized with TDP-43 and phosphorylated TDP-43, further supporting our postmortem tissue analysis and establishing a model system for future studies.

Overall, these observations align with a host of previous studies on N-protein, and for the first time, begin to link the cellular effects of N-protein and neurodegeneration. Previous *in vitro* studies have established that N-protein undergoes LLPS into BMCs (36–38). Using an optogenetic system, we characterized liquid-like biophysical properties of N-protein condensates. This study, however, extends our knowledge of LLPS dynamics in cells *in vitro*, and suggests that due to their observed low diffusivity, N-protein BMCs may form stable internal architectures (i.e., undergo LLPS maturation), as has been shown to occur with other RNA binding proteins (39, 40).

Our current study also aligns with previous research describing protein interactions with N-protein. Previous *in vitro* and *in silico* studies have described interactions of N-protein with TDP-43, FUS, hnRNP A1 and G3BP1 (25, 41–44). Like those studies, we support association of N-protein with TDP-43 and G3BP1, and to a lesser extent FUS and hnRNP A1, and further expand the repertoire of interacting proteins with N-protein. The differences between our observations for FUS and hnRNP A1 and those previously published may be due to the use of different experimental systems to study N-protein interactions (i.e., in cells *in vitro* compared to recombinant protein turbidity assays).

Current research suggests that the SARS-CoV-2 N-protein has transcriptional regulatory sequences that enhance the production of proteoform species, with their effect being to enhance SARS-CoV-2 viral fitness and suppress antiviral responses (26). In our current study, the proteoforms did not form detectable BMCs or colocalize with TDP-43 in our optogenetic assay. We begin to show and further theorize that BMC formation is dependent on aspects of the N terminal portion that are lost when the proteoforms are translated [i.e., post-translational modification sites, the Serine-Arginine (SR) domain, intramolecular amino acid interactions]. This postulate is strengthened by research showing that disruption of intramolecular amino acid interactions affects proper N-protein folding and RNA binding, and research demonstrating that phosphorylation within the SR domain affects N-protein oligomerization (45, 46).

A major field of interest in neurodegeneration, especially in ALS pathophysiology, is the dysregulation of RNA metabolism and splicing. A multitude of studies have shown that altered TDP-43 molecular dynamics, due to nuclear loss and formation of cytoplasmic inclusions, alter both RNA metabolism and splicing [reviewed in (33)]. Like those studies, we observed TDP-43 nuclear loss, cytoplasmic mislocalization, and alterations in RNA metabolism and splicing following optogenetically induced N-protein condensate induction and TDP-43 association.

We postulate that the alteration of TDP-43 homeostasis associated with neuronal N-protein expression may have an influence on the process of neurodegeneration and potentially influence disease trajectory (47, 48). If true, this postulate has implications for the impact of the SARS-CoV-2 infection that gave rise to COVID-19 given the peak age-distribution in the third through sixth decades, constituting a cohort of individuals that most likely to harbour a premanifest neurodegenerative process.

While undertaking this study, we understand that there are major limitations to exploring the causal link between SARS-CoV-2 and ALS. From a human spinal cord tissue perspective, samples were taken from post-mortem tissue, without prior medical records on their history of SARS-CoV-2 infection, and therefore we cannot comment at this stage whether the samples had an active infection at the time of sampling, or whether the observed N-protein immunoreactivity reflects a post-infection state. To partially address this limitation, we have begun to test human ALS spinal cord samples for the presence of other SARS-CoV-2 structural proteins (i.e., Spike, Envelope and Membrane) to provide additional evidence regarding viral exposure or persistence (data not shown). Based upon the N-protein BMCs *in vitro*, we cannot comment on the state of the structured condensates (i.e., liquid-like vs coalesced gel-like structures), nor whether they are still undergoing dissociation, but at a slower diffusivity. Further, while LLPS has previously been described for N-protein, there is an incomplete analysis of N-protein LLPS in cells *in vitro*, and more importantly, how N-protein LLPS may affect downstream mechanisms, especially those involved in neurodegeneration. While we have begun to address this, future studies will elaborate on the alteration in downstream mechanisms, and their global effects on neuron survival.

In summary, this study identifies SARS-CoV-2 N-protein immunoreactivity in a subset of human ALS spinal cord samples and demonstrates *in vitro* that optogenetically induced N-protein condensates associate with TDP-43 and are accompanied by molecular changes consistent with perturbed TDP-43 biology. These findings support further investigation into whether SARS-CoV-2 exposure can influence the natural history or trajectory of neurodegenerative disease.

## CRediT authorship contribution statement

**Joseph-Patrick W.E. Clarke:** Conceptualization, Data curation, Formal analysis, Investigation, Methodology, Supervision, Validation, Visualization, Writing – original draft, Writing – review & editing. **Alexandra Keating:** Formal analysis, Investigation, Visualization. **Veronica Noches:** Formal analysis, Investigation, Visualization. **Crystal McLellan:** Formal analysis, Investigation, Writing – review & editing. **Davide Dulcis:** Resources, Writing – review & editing. **John Ravits:** Resources, Writing – review & editing. **Michael J. Strong:** Conceptualization, Funding acquisition, Project administration, Resources, Supervision, Validation, Writing – review & editing.

## Acknowledgements

This work was supported by grants from the Wiley Family Foundation and Canadian Institutes of Health Research (SOP-160442). Human CNS tissues from UCSD were provided by the Target ALS Postmortem Tissue Core. Human lung tissues were kindly provided by Professor James Goldman, Columbia University. We acknowledge the participants and their families for their generous contributions, and the research teams at the contributing institutions for the collection and distribution of these materials. We gratefully thank Dr. Sean Cregan (Western University) for his expertise, guidance and utilization of his facilities to produce Opto-NCP lentiviral particles.

## Data availability

Requests for data, resources and reagents should be directed to the co-corresponding author Michael J. Strong.

## Materials and Methods

### Key Resources Table

Listed are the reagents/resources utilized in this study. Their purchase source and catalogue identifiers are included.

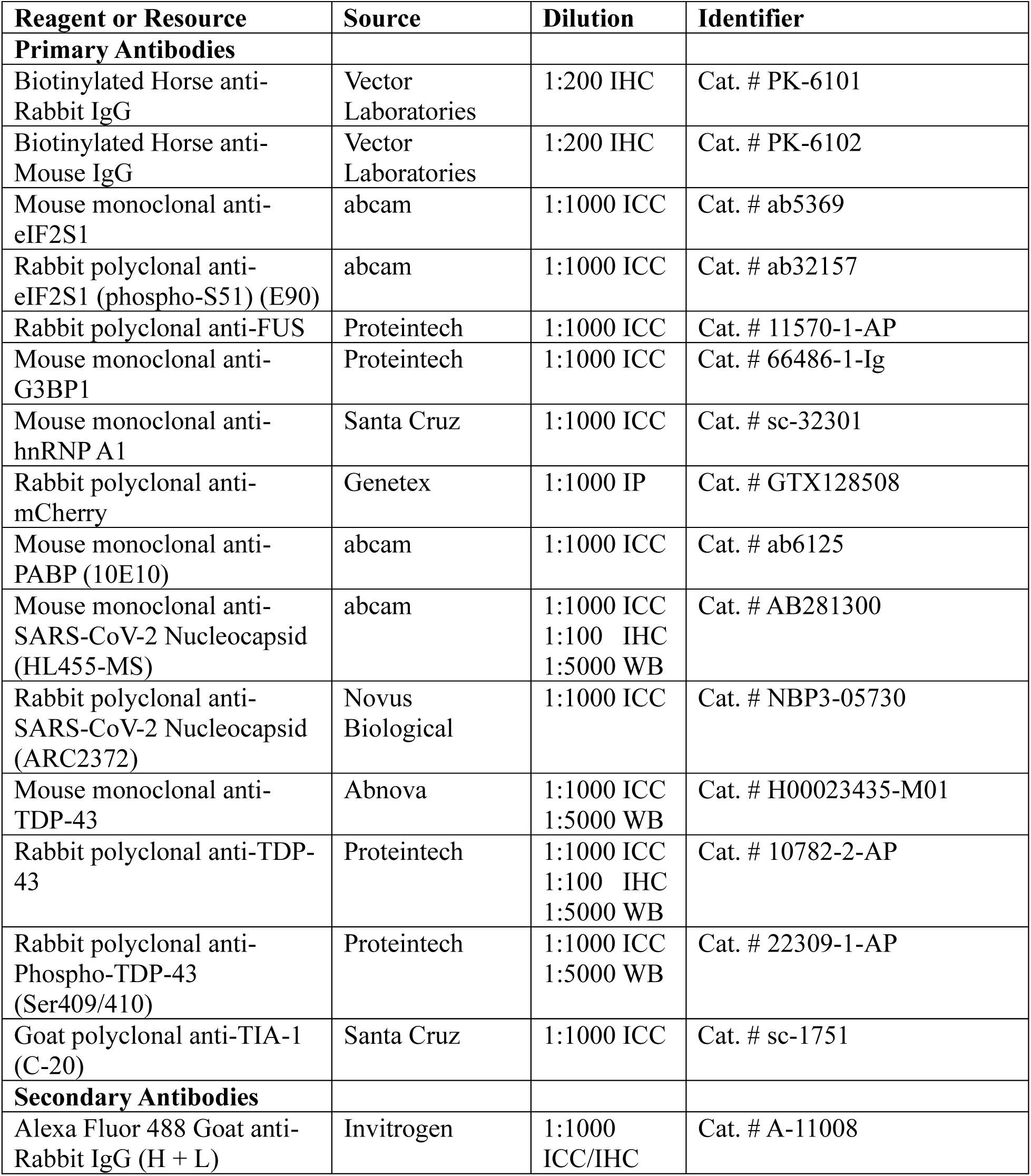
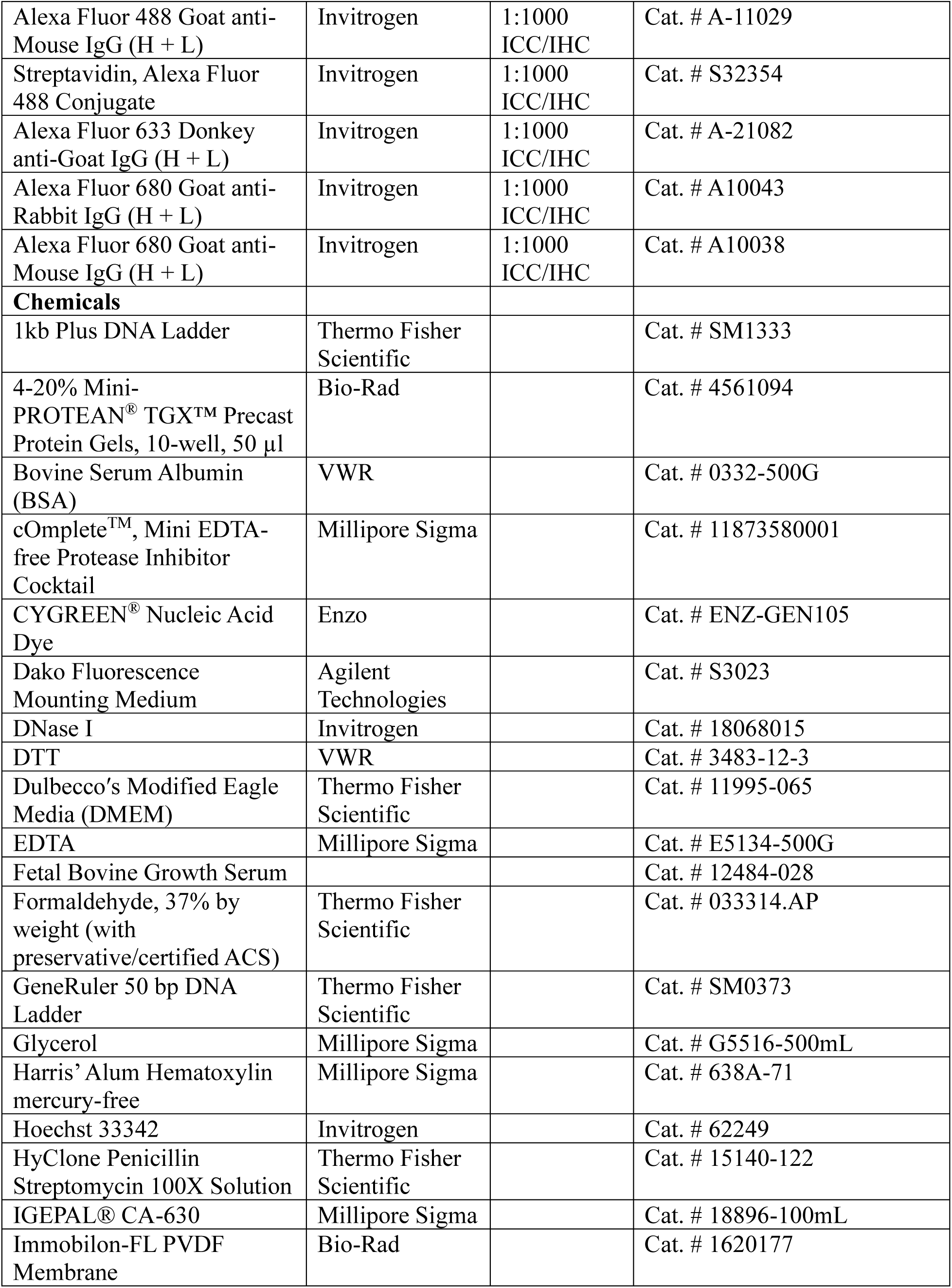
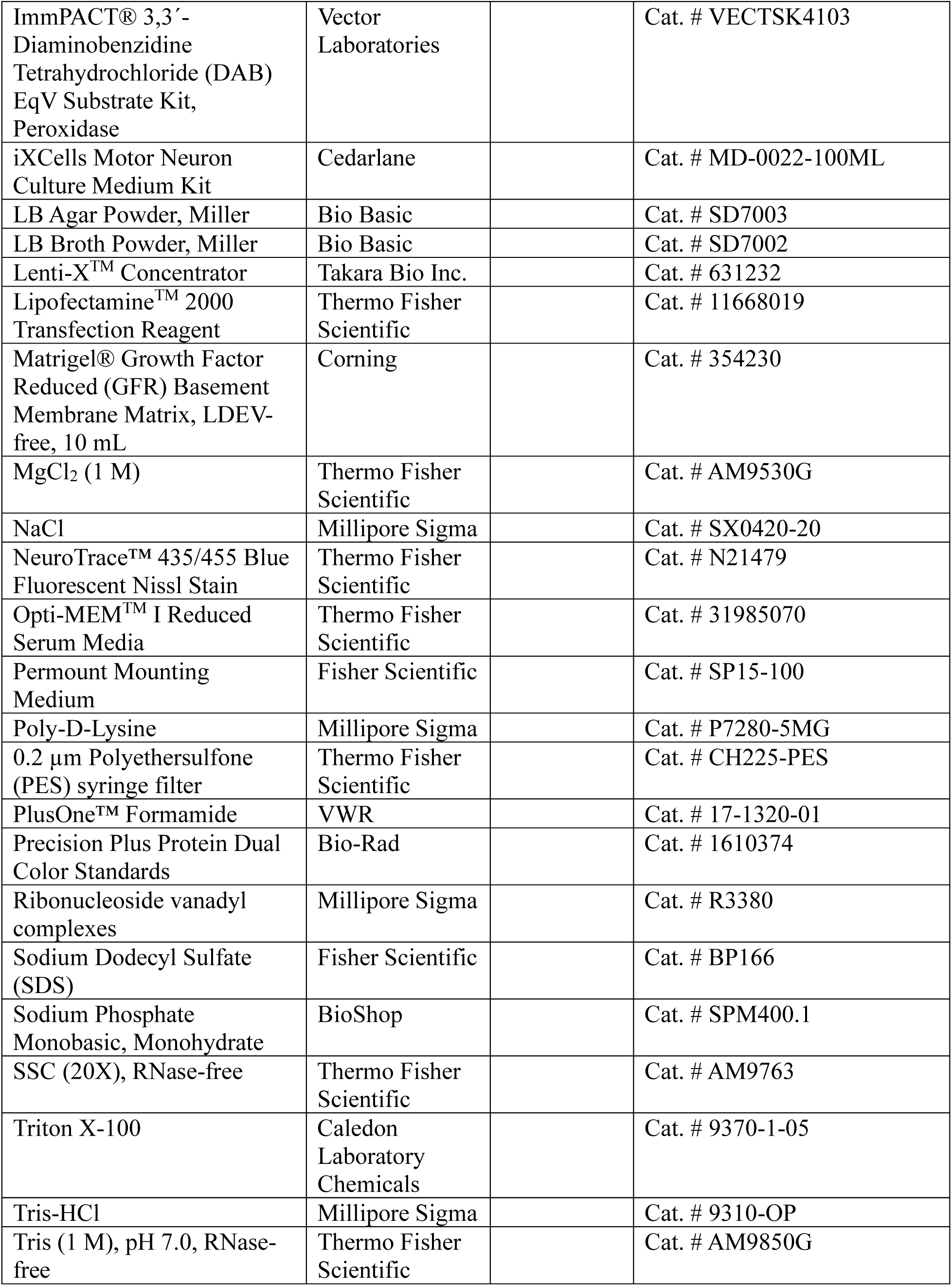
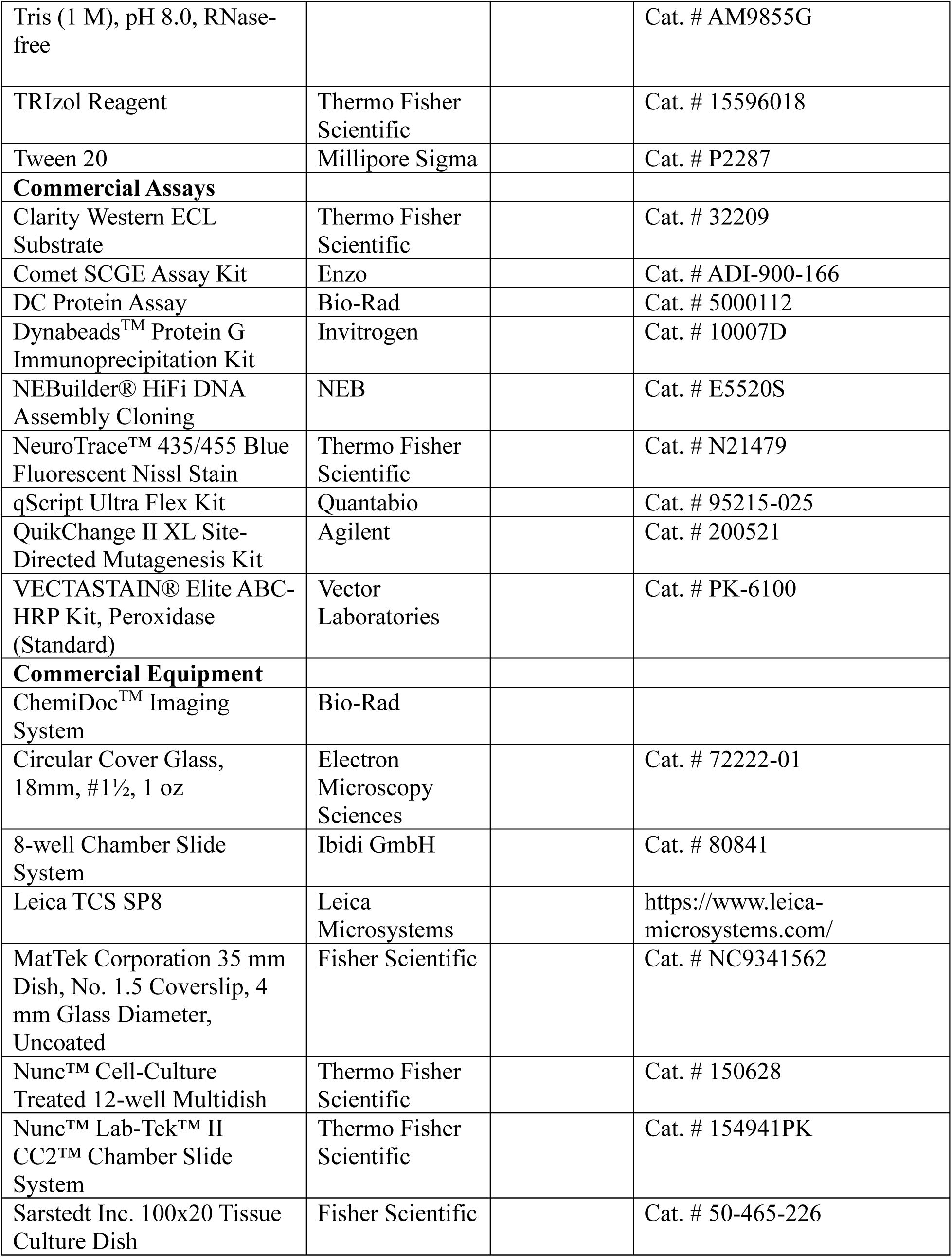
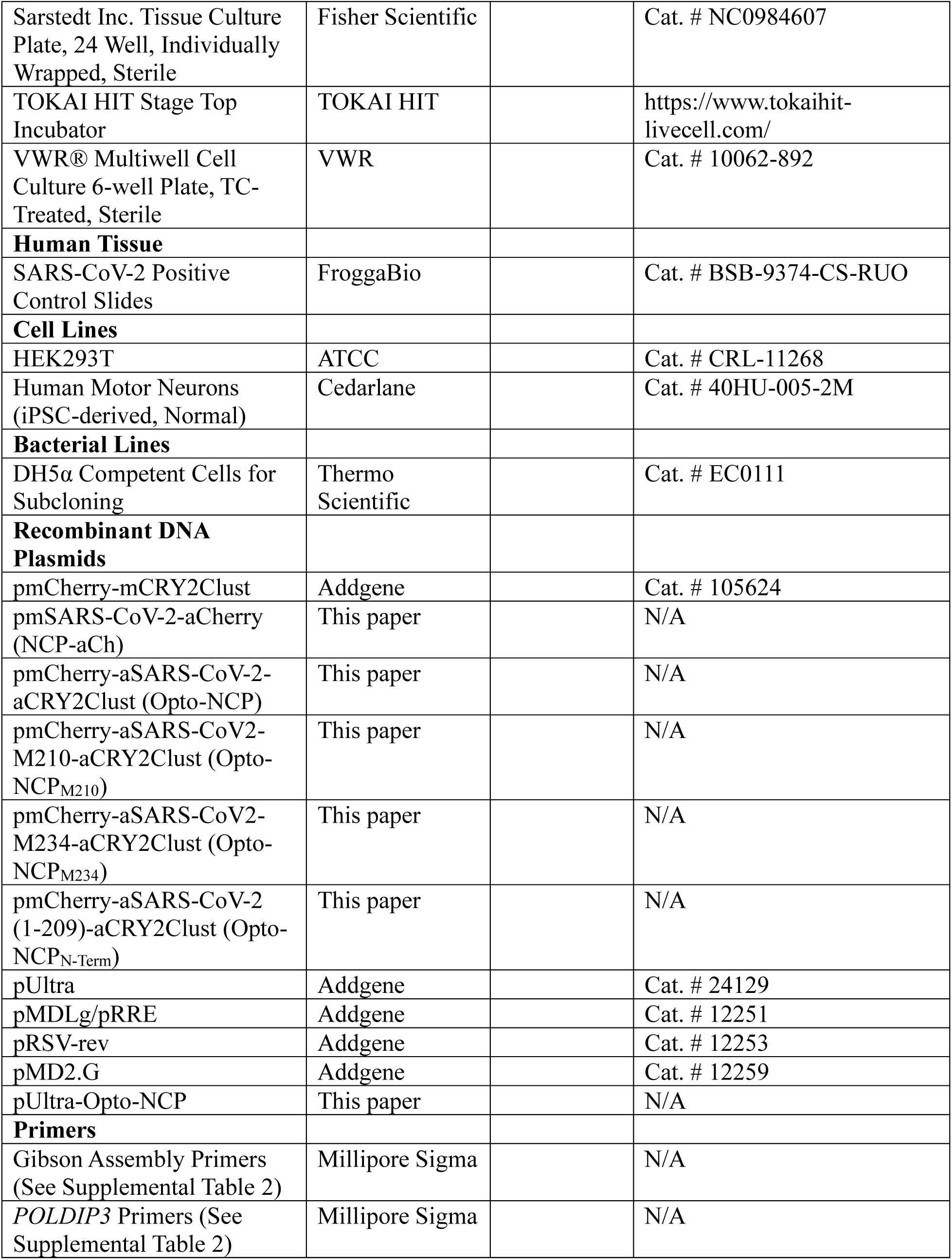
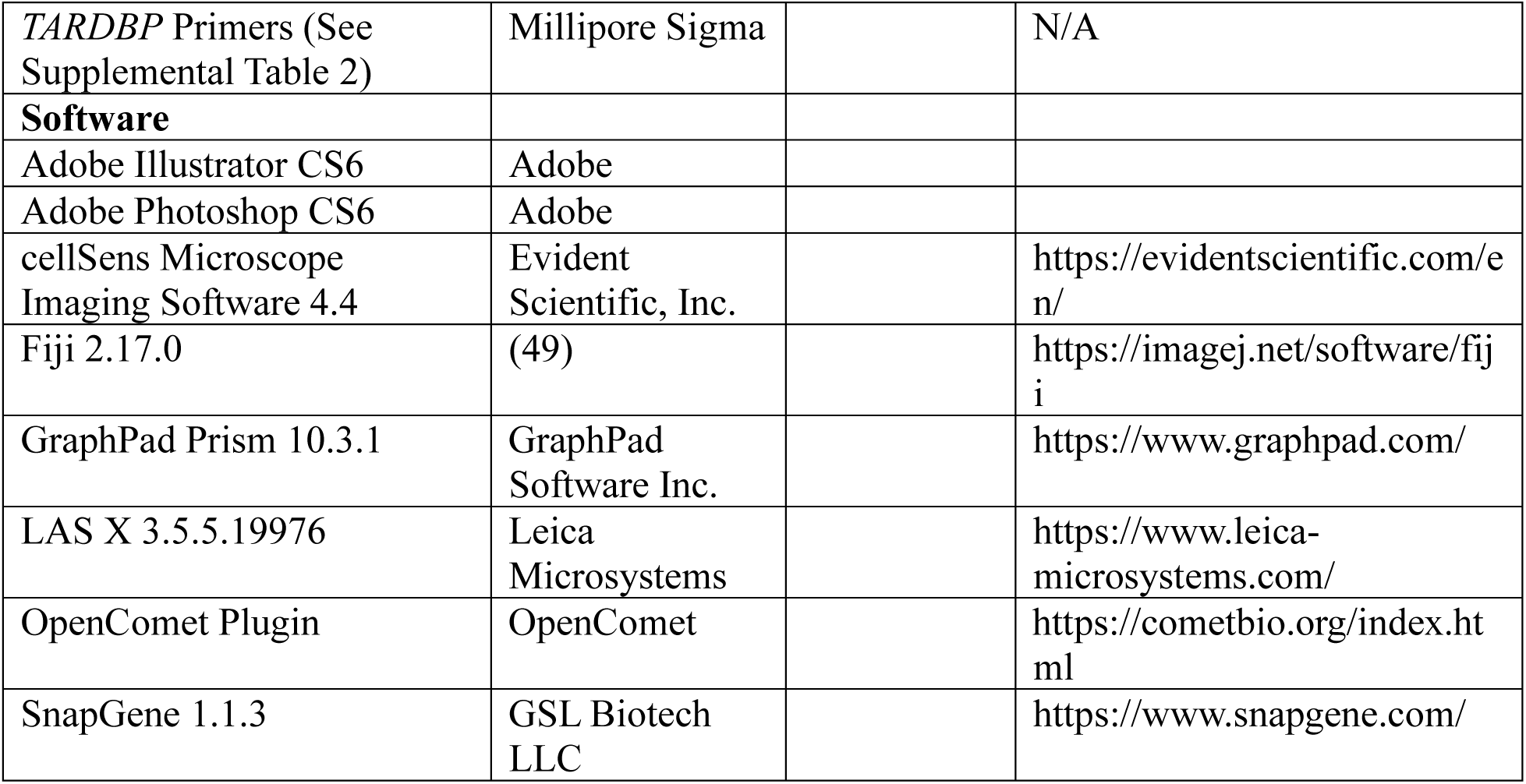

### Human Tissue

The ALS spinal cord cases (N = 36) used in this study were predominantly sporadic, with three familial cases also included (one SOD1 and two C9orf72). All samples were obtained between 2009 and 2022 at the University Hospital, London Health Sciences Centre (London, Ontario, Canada), using ethics research protocols approved by the Western University Health Sciences Research Ethics Board (HSREB) (HSREB #103735 & #124855). The study methodologies conformed to the standards set by the Declaration of Helsinki and its later amendments. Informed consent for autopsy and retention of tissues for research purposes was obtained in written format from all patients antemortem or, if not possible, from their spouse postmortem. No minors participated in this study. To ensure anonymization, at the time of utilization for this research project, all cases were assigned a unique identifier with the master list correlating the unique identifier maintained in a locked cabinet in the office of the principal investigator (Dr. Michael J. Strong) as per HSREB requirements.

Eight ALS lumbar spinal cord cases, collected during the SARS-CoV-2 pandemic, were from the ALS CNS Repository, Dr. John Ravits, University of California San Diego, California, USA through the Target ALS CNS Repository Consortium (https://targetals.org/research/human-postmortem-tissue-core/). Human tissues were obtained following HIPAA-compliant informed consent procedures approved by Institutional Review Board, University of California San Diego, San Diego, CA (IRB# 120056). Spinal cord tissues were acquired using a short-postmortem interval acquisition protocol usually under six hours. Tissues were immediately dissected in the autopsy suite, placed in labeled cassettes, and fixed in neutral buffered formalin for at least two weeks before being dissected and paraffin-embedded for indefinite storage.

SARS-CoV-2 positive control slides, containing SARS-CoV-2 infected human lung tissue samples, were obtained from FroggaBio, and were completely anonymous, as they were not provided with patient information nor contained any patient identifiers.

SARS-CoV-2 human lung tissue samples were obtained from Columbia University (a kind gift from Professor James Goldman, Columbia University, USA).

Supplemental Table 1 summarizes the demographics and disease features of the ALS patient tissues used in this study. The information is summarized from de-identified clinical and pathological records.

### Cell Culture

Human embryonic kidney 293T (HEK293T) cells were maintained in DMEM supplemented with 10 % fetal bovine serum and 1 % penicillin/streptomycin and incubated at 37 °C with 5 % CO_2_. SH-SY5Y cells were maintained in similar conditions. Cells were seeded into either 35 mm glass bottom imaging dishes, 12-well cell culture plates that were fitted with 18 mm circular glass coverslips, or 6-well culture plates. All imaging dishes and glass coverslips were coated with Poly-D-Lysine (50 mg/mL). Cell seeding was as follows: 50,000 per well in 35 mm glass bottom imaging dishes; 50,000 per well in 12-well culture plates; 150,000 per well in 6-well culture plates. 24 hours after seeding, cells were transfected with plasmid DNA, as outlined in the results section, using Lipofectamine 2000, as per the manufacturer’s protocol: 300 ng per well in 35 mm glass bottom imaging dishes; 300 ng per well in 12-well culture plates; 2.5 µg/well in 6-well culture plates. Cell cultures were protected from ambient light exposure and non-specific activation of the optogenetic protein, and all manipulations were performed in ambient light-free conditions. After overnight expression (16 hours) of the transfected plasmid DNA, experiments were conducted.

### Cloning and Site-Directed Mutagenesis

N-pmCherry-aSARS-CoV-2-N protein-aCRY2Clust-C (Opto-NCP) was constructed using full-length SARS-CoV-2 N protein (a gift from Dominic Esposito, Addgene, Plasmid # 170204) and N-pmCherry-mCRYClust-C (a gift from Won Do Heo, Addgene, Plasmid # 205624) using NEBuilder® HiFi DNA Assembly Cloning. To note, we positioned the CRY2Clust on the C-terminus of the N protein to distance it away from the N-terminal domain, so that it would A) not non-specifically influence N protein interaction dynamics, and B) allow for the assessment of N protein mutations for their influence on the overall kinetics of self-interaction. As a control for orientation, we generated a construct with the reverse orientation of Opto-NCP and observed similar BMC formation (data not shown). Briefly, CRY2Clust, N protein and mCherry were cloned into the base vector N-pmCRY2PHR-mCherry-C (a gift from Chandra Tucker, Addgene, Plasmid # 26866), replacing mCRY2PHR-mCherry, using the AfeI and NotI restriction enzyme sites of the backbone with four-fragment NEBuilder® HiFi Assembly. The methionine sites on CRY2Clust and N protein were mutated to alanine’s during cloning to avoid unwanted translation at these sites. A short flexible animo acid linker of GSSG (ggatcttctgga) was inserted during assembly between all three fragments to individualize all three fragments and both prohibit unwanted non-specific interactions and allow proper folding of each fragment.

As a non-Opto control, we also created N-pmNCP-aCherry-C. Briefly, NCP and mCherry were cloned into the into the base vector N-pmCRY2PHR-mCherry-C, replacing mCRY2PHR-mCherry, using the AfeI and NotI restriction enzyme sites of the backbone with two-fragment NEBuilder® HiFi Assembly Cloning. The methionine site on mCherry was mutated to alanine during cloning to avoid unwanted translation at that site. A short flexible animo acid linker of GSSG (ggatcttctgga) was inserted during assembly between both fragments to individualize both fragments and both prohibit unwanted non-specific interactions and allow proper folding of each fragment.

In this study, we also designed and tested N-pmCRY2Clust-aSARS-CoV-2-N protein (M210)-aCherry-C (Opto-NCP_M210_) and N-pmCRY2Clust-aSARS-CoV-2-N protein (M234)-aCherry-C (Opto-NCP_M234_). Briefly, truncated versions of the SARS-CoV-2-N protein (M210 and M234) were cloned into the base vector N-pmCRY2PHR-mCherry-C, replacing mCRY2PHR-mCherry, using the AfeI and NotI restriction enzyme sites of the backbone with three-fragment NEBuilder® HiFi Assembly. Given the absence of effect of reverse orientation with the Opto-NCP observed, this was not evaluated for truncations. The methionine sites on M210, M234 and mCherry were mutated to alanine’s during cloning to avoid unwanted translation at these sites. A short flexible animo acid linker of GSSG (ggatcttctgga) was inserted during assembly between all three fragments to individualize all three fragments and both prohibit unwanted non-specific interactions and allow proper folding of each fragment.

For this study, we also designed and tested the N-terminal fragment of N protein (amino acids 1-209), N-pmCherry-aSARS-CoV-2-N protein (N-Term)-aCRY2Clust-C (Opto-NCP_N-term_). Briefly, the N-terminus (including the N-arm and the N-terminal domain; amino acids 1-209) of the SARS-CoV-2-N protein was made via PCR from N-pmCherry-aSARS-CoV-2-N protein-aCRY2Clust-C and cloned back into N-pmCherry-aSARS-CoV-2-N protein-aCRY2Clust-C, replacing the full-length N protein gene, using BsrGI and KpnI restriction enzyme sites of the backbone with three-fragment NEBuilder® HiFi Assembly.

The expression of all transient expression constructs was driven by a CMV promoter. For recombinant protein production, however, a T7 promoter was used instead. Sequencing of the constructs was performed to confirm correct orientation, and to confirm the absence of unwanted mutations. All primer sequences used for cloning are listed in Supplemental Table 1.

### Lentivirus Cloning and Production

N-pmCherry-aSARS-CoV-2-N protein-aCRY2Clust-C (Opto-NCP) was subcloned into the pUltra 3^rd^ generation lentiviral vector (a gift from Malcolm Moore, Addgene, Plasmid # 24129) using the AgeI and EcoRI restriction enzyme binding sites of the backbone with two-fragment NEBuilder® HiFi DNA Assembly Cloning. Lentivirus generation was performed by first plating HEK293T cells in 10 cm TC-treated culture dishes and transfecting with a concoction of 10 µg pUltra-Opto-NCP, 9 µg pMDLg/pRRE (a gift from Didier Trono, Addgene, Plasmid # 12251), 4.5µg pRSV-rev (a gift from Didier Trono, Addgene, Plasmid # 12253) and 4.5µg pMD2.G (a gift from Didier Trono, Addgene, Plasmid # 12259). At 48 hours post-transfection, culture media was transferred to a 15 mL Eppendorf tube and centrifuged at 500 x g for 5 minutes to pellet cell debris. Cleared supernatant was the filtered through a 0.45µm Polyethersulfone (PES) filter. To the filtered supernatant, Lenti-X concentration solution was added at a ratio of 1:3.5 (Lenti-X:supernatant), mixed, and incubated at 4°C overnight. The next day, samples were centrifuged for 45 minutes at 4°C. The supernatant was carefully removed, and the viral pellets were resuspended in PBS (100 µL), aliquoted at 25 µL into pre-chilled 0.2 mL flat cap tubes and stored long-term at-80°C. To calculate virus titers, HEK293T cells were plated in 24-well plates and treated with a serial dilution of virus (10^-2^ to 10^-6^). After 48 hours, cells were visualized using an Olympus IX70 inverted phase contrast DIC fluorescence microscope, counting the fraction of mCherry positive cells at each virus dilution.

### Blue-Light (BL) Stimulation

BL stimulation was performed in Poly-D-Lysine (Millipore-Sigma) coated 35 mm glass bottom imaging dishes (live-cell imaging), 12-well plates fitted with 18 mm circular glass coverslips (fixed-cell imaging), or 6-well plates using custom-built LED arrays designed to fit multiple cell culture vessel dimensions and withstand common temperature/humidity requirements of cell culture incubators. LED arrays were positioned 5.0 cm above the culture surface, and 10,000 lux (measurement of luminous energy per unit of time, per unit area) of 465 nm BL was used to stimulate cultured cells.

### SDS-PAGE/Western Immunoblotting (WB)

Soluble proteins were extraction from HEK293T cells by first removing cell culture medium and replacing with ice-cold PBS. Cells were then scraped and collected in 1.5 mL Eppendorf tubes, followed by centrifugation at 500 x g for 5 minutes at 4°C. PBS was aspirated and cell pellets were resuspended in lysis buffer (25mM Tris-HCl, pH 7.4; 150mM NaCl; 1mM EDTA; 1% IGEPAL® CA630; 5% glycerol; 1 cOmplete, Mini, EDTA-free Protease Inhibitor Cocktail tablet). The cell suspension was then sonicated (2 × 10 second pulses on ice) and pre-cleared via centrifugation (16,000 x g for 5 minutes at 4°C). Protein concentrations were determined using the DC Protein Assay, as per the manufacturer’s protocol. Equivalent amounts of total protein were mixed with 5X Laemmli buffer, supplemented with DTT, and denatured at 95°C for 5 minutes. Protein separation was performed using 4-20% SDS-PAGE and transferred to PVDF membranes using the wet transfer method. Ponceau S staining was used to assess the equivalency of protein loading after SDS-PAGE transfer. Membranes were washed with 1.0M Tris-buffered saline (TBS), pH 7.4, blocked with 5% bovine serum albumin (BSA) in TBS for 1 hour at room temperature, and then incubated with primary antibodies (Key Resources Table; all primary antibodies were used at a concentration of 1:1000) in 5% BSA in TBS-T (0.1% Tween 20) overnight at 4°C. Following TBS-T washes, membranes were incubated with peroxidase-conjugated secondary antibodies in TBS-T (Key Resources Table; all secondary antibodies were used at a concentration of 1:3000) and protein bands were detected using Clarity Western ECL Substrate and visualized using the ChemiDoc XRS+ Gel Imaging System.

For sequential probing, membranes were incubated with 30% H_2_O_2_ for 15 minutes at room temperature, followed by a 5-minute TBS wash. This completely inactivates the secondary antibody HRP bound to the first probed antibody and does not require a blot stripping step. Membranes were then re-blocked in 5% BSA in TBS for 1 hour at room temperature, followed by incubation with new primary antibodies in 5% BSA in TBS-T overnight at 4°C. To note, this was only performed when using sequential antibodies raised in different hosts (i.e., Rabbit antibody followed by Mouse antibody, or vice versa).

### Co-Immunoprecipitation

Soluble proteins were extracted from HEK293T cells, as described above. Immunoprecipitation was performed using the Dynabeads^TM^ Protein G Immunoprecipitation Kit, as per the manufacturer’s protocol with minor experimental modifications. Briefly, antibody binding used 2.2µg of rabbit polyclonal anti-mCherry antibody for each immunoprecipitation, and the antibody-dynabead mix was incubated with rotation for three hours at room temperature. Immunoprecipitation of the target antigen used 500 µg of total cell lysate and was incubated with rotation for four hours at 4°C. After isolation and washing, the dynabead antibody-antigen complexes were resuspended in 5X Laemmli buffer, supplemented with DTT, and denatured at 95°C for 5 minutes. Protein separation was performed using 4-20% SDS-PAGE and transferred to PVDF membranes using the wet transfer method. Further probing and development of membranes were like those outlined above.

### Immunoprecipitation Mass Spectrometry (IP-MS)

HEK293T cells were transfected with our Opto-NCP plasmid (see above), left to express overnight, and then subjected to either BL stimulation, or left in the dark, for 60 minutes (see above). In parallel, HEK293T cells that were non-transfected, and not subjugated to BL stimulation were also grown. Immediately after BL stimulation, soluble protein extracts for each condition were acquired, and underwent immunoprecipitation until the elution phase of the protocol. After samples were eluted from the Protein G Dynabeads^TM^, they were frozen at-80°C and then sent to Creative Proteomics (Creative Proteomics, New York, USA) for mass spectrometry analysis, as per their protocol. Briefly, once obtained, samples were run on a 12% SDS-PAGE and silver stained to assess the quality of each sample. Next, the lanes containing protein were cut into 1 mm^3^ cubes and transferred to 1.5 mL microcentrifuge tubes. To the samples, 1.0 mL of a 50/50 mix of 30 mM K_3_Fe(CN)_6_ and 100 mM Na_2_S_2_O_3_ was added. When the brown colour of the silver stain disappeared, the supernatant was removed and the reaction was stopped by adding 200 µL of ultrapure water. After 10 minutes, the supernatant was discarded and 1.0 mL of ammonium bicarbonate was added for 30 minutes. Again, the supernatant was discarded and 500 µL of acetonitrile was added for 30 minutes. The acetonitrile was then removed, and the gel slice was rehydrated in 10 mM DTT and incubated at 56°C for 60 minutes. After heating, the DTT was removed and the gel slice was incubated for 10 minutes at room temperature with 500 μL of acetonitrile. The supernatant was discarded and the gel slice was incubated for 30 minutes at room temperature in the dark with 50 mM iodoacetamide. After 30 minutes, the supernatant was removed and the gel slice was incubated with 500 µL of acetonitrile for 10 minutes at room temperature. After incubation, the supernatant was removed and trypsin was added to the gel slice with incubation on ice for 45 minutes. After 45 minutes on ice, the sample was incubated overnight at 37°C. The supernatant was then transferred to a fresh 1.5 mL microcentrifuge tube, to which a 50 mM ammonium bicarbonate/acetonitrile solution (1:2, v/v) was added and incubated for 60 minutes at 37°C. The solution was then transferred to a fresh 1.5 mL microcentrifuge tube, and 0.1% trifluoroacetic acid was added and incubated for 30 minutes. After incubation, 50 mM ammonium bicarbonate/acetonitrile solution (1:2, v/v) was added. The extracts were then combined and lyophilized to near dryness. The samples were then resuspended in 20 µL of 0.1% formic acid before nanoscale liquid chromatography coupled to tandem mass spectrometry (Nano LC-MS/MS) analysis.

The Nano LC-MS/MS was run using an Ultimate 3000 nano UHPLC system coupled with a Q Exactive HF mass spectrometer with an ESI nanospray source. The nanocolumns used were the PepMap C18, 100Å, 100 μm × 2 cm, 5.0 μm (trapping column) and the PepMap C18, 100Å, 75 μm × 50 cm, 2.0 μm (analytical column). The full scan for mass spectrometry was performed between 300–1,650 m/z at the resolution 60,000 at 200 m/z, the automatic gain control target for the full scan was set to 3e6. The MS/MS scan was operated in Top 20 mode using the following settings: resolution 15,000 at 200 m/z; automatic gain control target 1e5; maximum injection time 19ms; normalized collision energy at 28%; isolation window of 1.4 Th; charge sate exclusion: unassigned, 1, > 6; dynamic exclusion 30 seconds.

### Immunocytochemistry (ICC)

Circular glass coverslips seeded with cells were prepared for ICC by fixation with 3.7 % formaldehyde in 1.0 M PBS, pH 7.4, for 15 minutes at room temperature. Following fixation, cells were washed three times with PBS and were permeabilized with 0.3 % Triton X-100 in PBS for 30 minutes. Three more washes were performed with PBS, and the cells were then blocked with 5% BSA in PBS for 60 minutes at room temperature. Cells were then washed three times with PBS and were incubated overnight at 4 °C with primary antibodies (Key Resources Table) diluted in PBS with 5 % BSA. The next day, the cells were washed three times with PBS and then were incubated with secondary antibodies (Key Resources Table) diluted in PBS with 5 % BSA for 60 minutes at room temperature. Cells were then washed three times with PBS and incubated with Hoechst 33342 (1.0 µg/mL diluted in PBS) for 5 minutes at room temperature, followed by washing with PBS. Finally, the circular glass coverslips were mounted onto microscope slides with Dako Fluorescence Mounting Medium. All images and quantifications are representative of three biological replicates. Fixed-cell imaging was performed using a Leica TCS SP8, inverted, confocal microscope using the LAS X software. 63X random fields of view (Leica high contrast/high colour, plan apochromat 63X/1.40 Oil, coverslip series 2 objective: 3 random fields per experiment) were acquired for quantification, and approximately 50 cells were analyzed per experiment, per condition, per replicate. All images were acquired using identical imaging settings within each experiment. Briefly, on average the laser intensity of the Hoechst, 488nm and mCherry channels were: 1%, 0.1% and 0.5%, respectively. Each image was z-stack imaged at 0.3 µm/stack, with, on average, 30 z-stacks taken per image. Images were then post-processed using the LAS X Lightning Process (Refractive Index: 1.372) to eliminate background signal and resolve nanostructures as small as 120 nm. Final images are max projections exported from the LAS X software.

### Immunohistochemistry (IHC)

#### 3,3‘-Diaminobenzidine Tetrahydrochloride (DAB)

DAB immunohistochemistry was completed on paraffin embedded, formalin fixed postmortem human cervical spinal cord tissue. Briefly, glass slides with cut tissue (10µm slices) were deparaffinized on a slide warmer set to 55°C for 15 minutes. Deparaffinized slides were then rehydrated with three 5-minute washes of xylene and successive 5-minute washes of ethanol at 100%, 95% and 80%, followed by a final deionized water wash. Slides were then placed in a plastic boat container with sodium citrate buffer (10mM sodium citrate, 0.05% Tween 20, pH 6.0) and underwent antigen retrieval using a pressure cooker (2,100 Retriever; Aptum Biologics) heated to 120°C for 30 minutes. After slides were removed and cooled to room temperature, they were quenched in 3% hydrogen peroxide for 15 minutes at room temperature and subsequently washed in PBS for 15 minutes. Slides were then blocked (5% bovine serum albumin, 0.3% Triton-X 100 in PBS) for one hour at room temperature. Primary antibodies were then added (Key Resources Table), diluted in blocking solution, and left to incubate in a humified chamber overnight at 4°C. The following day, slides were washed three times with PBS and biotinylated secondary antibodies, diluted in blocking solution, were added and incubated for one hour at room temperature. Depending on the primary antibodies used, either a biotinylated horse anti-mouse IgG secondary antibody or a biotinylated goat-anti-rabbit IgG secondary antibody was used on the slides. The slides were treated with the VECTASTAIN Elite ABC peroxidase system, per the manufacturer’s protocol, followed by saturation with DAB (saturation time was determined microscopically for each individual antibody), and a counterstain using Harris’ Alum Hematoxylin mercury-free for 25 seconds. Water, ethanol and xylene washes, in opposite order from rehydration performed above, were performed before the slides were mounted using Permount mounting medium. Slides were visualized using an upright Olympus BX45 pathology microscope, fitted with an Olympus CAM-SC180, colour, CMOS, camera. Images of the alpha motor neurons in the anterior horns of the cervical spinal cord tissue were captured using either an Olympus Plan N 40X/0.65 Ph2 or an Olympus Plan N 100X.1.25 Oil objective. Further image processing was performed using cellSens microscope imaging software.

#### Tissue Confocal Immunofluorescence (Confocal IF)

Tissue confocal immunofluorescence was completed on paraffin embedded, formalin fixed postmortem human cervical spinal cord tissue. Slides were initially UV (380nm) treated overnight. After UV treatment, slides were developed similarly to immunohistochemistry slides (deparaffinization, rehydration, antigen retrieval and serum blocking) (see above methodology). Primary antibodies were added, diluted in blocking solution, and left to incubate in a humified chamber overnight at 4°C. The following day, slides were washed three times with PBS and fluorescent secondary antibodies, diluted in blocking solution, were added and incubated for one hour at room temperature. Slides were then counterstained using NeuroTrace^TM^ 435/455 Blue fluorescent Nissl stain, as per the manufacturer’s protocol. Slides were then mounted using Dako Fluorescence Mounting Medium. Fixed-tissue imaging was performed using a Leica TCS SP8, inverted, confocal microscope using the LAS X software. Images of the alpha motor neurons in the anterior horns of the cervical spinal cord tissue were acquired using a Leica high contrast/high colour, plan apochromat 63X/1.40 Oil, coverslip series 2 objective. All images were acquired using identical imaging settings within each experiment. Briefly, on average the laser intensity of the NeuroTrace^TM^ 435/455 Blue fluorescent Nissl stain, 488nm and 680nm channels were: 0.1%, 0.05% and 0.25%, respectively. Each image was z-stack imaged at 0.3 µm/stack, with, on average, 30 z-stacks taken per image. Images were then post-processed using the LAS X Lightning Process (Refractive Index: 1.372) to eliminate background signal and resolve nanostructures as small as 120 nm. Final images are max projections exported from the LAS X software.

RNA Extraction and First-Strand Complementary DNA Synthesis and RT-PCR for Measuring *TARDBP* and *POLDIP3* Expression and Splice Variants

RNA from cell samples was isolated using the TRIzol reagent, as per the manufacturer’s protocol. RNA concentrations were determined using a NanoDrop^TM^ One Microvolume UV-VIS Spectrophotometer. 1.0 µg of isolated RNA was further DNase I treated, as per the manufacturer’s protocol. First-strand cDNA was synthesized from RNA samples using the qScript® Ultra Flex Kit, as per the manufacturer’s standard reaction protocol with minor experimental modifications. Briefly, 100ng of DNase I treated RNA was used for each reaction (25 µL), where gene-specific reverse primers for *TARDBP* or *POLDIP3* were used instead of Oligo(dT) or random primers (See Supplemental Table 1). The following cycling parameters were used: 98°C/30 seconds, 40 cycles @ (98°C/10 seconds, 62°C/15 seconds, 72°C/30 seconds), 72°C/2 minutes, 4°C/hold. The PCR products were visualized on a 2% agarose TAE gel using a 50 bp ladder.

### CometAssay®

The CometAssay® was conducted using a single-cell gel electrophoresis (SCGE) assay kit, following the manufacturer’s protocol. Briefly, cells were plated in 6-well plates and transfected with either NCP-aCh or Opto-NCP. 24 hours after transfection, cells were treated with or without BL stimulation for 360 minutes. Additionally, a separate plate of cells was treated with methylglyoxal (400 µM; cell stress control) for 24 hours. After treatments, the cells were collected, washed with ice-cold PBS, and resuspended in ice-cold PBS (without Mg^2+^ or Ca^2+^) at a final concentration of 1 × 10^5^ cells/mL. The cells were then mixed with low-melting-point agarose at a volume ratio of 1:50, and 75μL aliquots were loaded onto slides. The slides were incubated in a dark and humid environment at 4°C for 30 minutes, followed by overnight incubation in a pre-chilled lysis solution from the kit. The next day, the slides were placed in an alkaline solution (pH > 13) for 60 minutes at room temperature, in the dark. Electrophoresis of the slides was performed at 300 mA in alkaline buffer for 40 minutes under dark conditions at 4°C. The slides were then washed in water, fixed in 70% ethanol for 5 minutes, and allowed to dry completely. DNA was then stained with CYGREEN dye for 30 minutes. The slides were examined using a Leica TCS SP8, inverted, confocal microscope using the LAS X software. At least 100 individual cells per sample were evaluated in quadruplicate using Fiji (OpenComet plugin).

### Live-Cell Imaging

Live-cell imaging was performed using a Leica TCS SP8, inverted, confocal microscope mounted with a TOKAI HIT heating, humidity and CO_2_ chamber that was regulated by a TOKIA HIT STX temperature and CO_2_ module. Additionally, the objective in use for live-cell imaging was wrapped with a TOKAI HIT lens heater. 35 mm glass bottom imaging dish were equilibrated on the microscope at 37 °C and 5 % CO_2_ for 15 minutes prior to imaging. BL stimulation was performed using the custom-built LED array positioned above the TOKAI HIT chamber to stimulate cells in the live-cell incubation unit (5.0 cm above the glass culture surface). Cells were stimulated for 60 minutes (465 nm, 10,000 lx) to induce Opto-NCP biomolecular condensate formation. For imaging, we captured using a Leica high contrast/high colour, plan apochromat 40X/1.30 Oil, coverslip series 2 objective, in 10-minute intervals over the total 60 minutes to account for thermal drift during imaging. The 10-minute intervals were programmed using the FRAP module in LAS X, where we set the FRAP intensity to zero percent to allow for non-FRAP stimulated live-cell imaging. During the 10-minute stimulation intervals, the BL LED was intermittently turned on for 11 seconds and turned off for 11 seconds to allow for the LAS X microscope software to capture the mCherry signal in the 594 nm channel without stimulation occurring during image capture. These intervals were achieved by synchronizing the LED and FRAP module using a Nearpow Multifunctional Infinite Cycle Programmable Plug-in Digital Timer Switch (Nearpow) attached to and controlling the LED array. Data is representative of three biological replicates, with approximately 50 cells analyzed per condition, per replicate.

### Fluorescence Recovery After Photobleaching (FRAP)

FRAP was performed using a Leica TCS SP8, inverted, confocal microscope mounted with a TOKAI HIT heating, humidity and CO_2_ chamber that was regulated by a TOKIA HIT STX temperature and CO_2_ module. Additionally, the objective in use for live-cell imaging was wrapped with a TOKAI HIT lens heater. 35 mm glass bottom imaging dish were equilibrated on the microscope at 37 °C and 5 % CO_2_ for 15 minutes prior to imaging. BL stimulation was performed using the custom-built LED array positioned above the TOKAI HIT chamber to stimulate cells in the live-cell incubation unit (5.0 cm above the glass culture surface). Cells were stimulated for 60 minutes (465 nm, 10,000 lx) to induce Opto-NCP biomolecular condensate formation. Images were captured using a Leica high contrast/high colour, plan apochromat 40X/1.30 Oil, coverslip series 2 objective, in 10-minute intervals to account for thermal drift during imaging. The 10-minute intervals were programmed using the FRAP module in LAS X, where we set the FRAP intensity to zero percent to allow for non-FRAP stimulated live-cell imaging. For FRAP analysis, after BL stimulation we then re-calibrated the FRAP module in LAS X to induce a 20% laser bleach for 1.7 seconds in a circular 2.0 µm^2^ area partially encompassing an Opto-NCP biomolecular condensate. In this manner, we were able to assess aspects of diffusion and mobility of the Opto-NCP biomolecular condensates, both within the biomolecular condensates themselves and with the free Opto-NCP within the cytoplasmic environment. FRAP data was analyzed during the 90 seconds post-bleach, and is representative of three biological replicates, with approximately 10 biomolecular condensates analyzed per replicate.

### iPSC Human Motor Neuron Cultures

For iPSC human motor neuron cultures, chamber slides (8-well, Nunc™ Lab-Tek™ II CC2™) were first coated with Matrigel GFR basement membrane matrix, as per the manufacturer’s protocol. To note, the chamber slides used in this protocol feature a chemically modified CC^2^ growth surface that mimics the characteristics of poly-D-lysine coating. We also used 8-well ibidi GmbH chamber slides that we coated with poly-d-lysine at 10.0 µg/mL prior to Matrigel coating. iPSC human motor neurons were handled as per the manufacturer’s protocol. Briefly, iPSC human motor neurons (derived from a healthy 24-year-old Caucasian male) were thawed, seeded at 100,000 cells per well and maintained in iXCells motor neuron culture medium. Half-media changes were performed every two days for seven days, at which time the cultures were fully differentiated into human motor neurons, expressing high level of ChAT and MAP2. Differentiated human motor neurons were then transduced with Opto-NCP lentivirus (see above for methodology), measuring expression after 48 hours. Once expressing Opto-NCP, the motor neuron cultures underwent BL experimentation and analysis.

### Quantification and Statistical Analysis

#### Immunocytochemistry

Image visualization and cell counting of Opto-NCP transfected, non-cluster and Opto-NCP-cluster containing cells, were performed with LAS X software. *Z*-Stack images were taken at 0.3μm slices with 30 z-stacks per image on average. Images were post-processed using the LAS X Lightning Process (Refractive Index: 1.372) to eliminate background signal and resolve nanostructures as small as 120 nm. Final images are max projections exported from the LAS X software as 16-bit.TIFF images.

#### Immunoprecipitation Mass Spectrometry (IP-MS)

The raw mass spectrometry files were analyzed and searched against *Homo sapiens* protein database according to sample species using Maxquant (1.6.2.6). The parameters were set as follows: the protein modifications were carbamidomethylation (C) (fixed), oxidation (M) (variable); the enzyme specificity was set to trypsin; the maximum missed cleavages were set to 2; the precursor ion mass tolerance was set to 10 ppm, and MS/MS tolerance was 0.05 Da. For statistical comparisons of results, a one-tailed, two-sample unequal variance, t-Test was performed between non-stimulated and BL stimulated sample results (*p* < 0.05, C.I. = 95%). The *p*-values from the t-Test’s were then converted to-log10 (*p*-value) for easier scaling and analysis on a Volcano plot [*p*-value 0.05 =-log10 (*p*-value) 1.30]. The fold change (FC) between samples was calculated by dividing the average BL stimulated sample intensity by the average non-stimulated sample intensity. For easier scaling and analysis on a Volcano plot, the fold change was converted to log_2_ (FC), where values ≥ 1 are considered increased [log_2_ (FC) 1 = twofold increase; doubling], and ≤-1 are considered decreased [log_2_ (FC) 1 = twofold decrease; halving].

#### Opto-NCP BMC Analysis

Images were further processed using the Fiji fast Fourier transform (FFT) bandpass filter (larger structures down filtered to 40 pixels; small structures filtered up to 1 pixel) to smooth out small structures for analysis. Opto-NCP BMCs were analyzed by thresholding individual cytoplasmic BMCs and using the Analyze Particles tool in Fiji. Output data gave the area (μm^2^) and perimeter (μm) of each individual BMC. The average number of BMCs per cell was calculated by dividing the total number of BMCs counted by the by number of cells counted, per experiment, and then averaged over three replicate experiments.

#### Colocalization

##### ICC

Fixed-cell image quantification of Opto-NCP biomolecular condensates was examined using Fiji, with Pearson’s correlation coefficient for Opto-NCP biomolecular condensate and RNA binding protein (RBP) co-localization calculated using the BioImaging and Optics Platform (BIOP), with approximately 5 fields of view analyzed per condition, encompassing approximately 50 cells and 200 clusters analyzed over triplicate experiments. Briefly,.LIF files from LAS X were open in Fiji, with individual z-stacked files chosen for analysis. The z-stacked files were made into multichannel composite images, and individual biomolecular condensates were selected as ROIs using the freehand selection tool. The ROIs were then run through the BIOP, using the Just Another Colocalization plugin (JaCoP) in the Image Analysis folder. In the tooltip, the green and red channels were chosen for colocalization (Green = RBP, Red = Opto-NCP), images were automatically thresholded using Otsu, and z-slices were considered separately to calculate Pearson’s correlation coefficients. Only z-slices within a given ROI that contained both channels were considered for colocalization analysis. If a ROI had multiple z-slices that exhibited zero or positive Pearson’s correlation coefficients, these were averaged to obtain an overall Pearson’s correlation coefficient for the ROI.

##### IHC

NCP biomolecular condensates were defined based upon the presence of multiple densely compact deposits within the cytoplasm, and with the absence of any nuclear expression. For quantification of NCP biomolecular condensate and TDP-43 colocalization in post-mortem motor neurons, a similar approach was taken as outlined above.

#### Protein Expression Analysis Based on Fluorescence

Cells containing Opto-NCP clusters were outlined and fluorescence measurements of eIF2S1 and phospho-eIF2S1 were obtained. Output fluorescence data gave the integrated density (*i.e.*, the sum of the values of the pixels in the image or selection) of fluorescence. These measurements were then background corrected using the Correct Total Cell Fluorescence (CTCF) formula: *CTCF = Integrated Density – (Area of Selected Cell x Mean Fluorescence of Background Readings)*

For background correction, the image background fluorescence was determined from the average of three random regions of interest that contained no fluorescent signal on the image analyzed. The CTCF of all analyses was compared to the no BL stimulation and no treatment group and represented as a fold change. Overall, approximately 50 cells were analyzed per condition per biological replicate. Analysis scales were set based upon the magnification of the images taken.

#### Live-Cell Imaging – Association and Dissociation Kinetics

Images from every 10-min time-point in the live-cell experiment were exported as 16-bit.TIFF images from LAS X. Exported images were imported into Fiji and converted to 8-bit images. Analysis of cells containing Opto-NCP clusters was performed manually. Specifically, the number of cells containing detectable Opto-NCP clusters were tracked over time and divided by the total number of cells within the imaging field to generate a percentage of cell with inclusions for each time-point within the imaging sequence. For Opto-NCP cluster association and dissociation rate quantification, cells with clusters were initially counted for each time-point. Each time-point cluster value was then converted to a percentage of the total cellular cluster maximum at the height of the stimulation cycle. Mean percentages for each time-point, from three biological replicates, were then plotted over-time for each stimulation cycle. The kinetic rate of association and/or dissociation is noted as the half-maximal kinetic association/dissociation rate (KA_1/2Max_ or KD_1/2Max_, respectively).

#### FRAP

FRAP quantification of Opto-NCP biomolecular condensates was examined using Fiji. Briefly,.LIF files from LAS X were opened in Fiji. Time-lapses of individual biomolecular condensates that underwent FRAP were cropped using the freehand selection tool and duplicated. For analysis, the individual cropped time-lapses were run using plugins from the “Stowers ◊ Jay_Unruh ◊ Image Tools” folder. Specifically, each BMC time-lapse was first run using the “create spectrum jru v1” plugin, selecting the “average” spectrum statistic. These were saved as individual “Plot Objects”. Next, each time-lapse “Plot Object” was opened, and combined using the “combine all trajectories jru v1” plugin. After combining all BMC “Plot Objects”, the data was normalized using the “normalize trajectories jru v1” plugin, selecting the “Min_Max” normalization. The normalized data was then run using the “batch FRAP fit jru v1” plugin, manually selecting the “Frames before FRAP”, “Frames to analyze” and “Maximum tau (frames)”, with the data output saved and opened in Excel. Additionally, the “List” data from the analysis was saved and opened in Excel for further graphical analysis in GraphPad Prism.

#### Statistics

Statistical significance was calculated in GraphPad Prism software (Version 10.3.1) and resulting *p* values less than or equal to 0.05 [95% Confidence Interval (CI)] were determined to be significant. One-way ANOVAs, with Tukey post-hoc analysis, and two-way ANOVAs, with Bonferroni post-hoc analysis, were used where indicated. To determine kinetic, half-maximal association and dissociation responses, graphs were subjected to a non-linear regression curve fit, using an (agonist) vs normalized response with a variable slope least squares fit equation. *n* values described in the text and figure legends represent number of cells per experimental group, unless otherwise indicated, across multiple independent, biological experiments. Representative images and videos were prepared using Adobe Illustrator CS6 and Photoshop CS6 (Adobe).

## Supplemental Figure Legends

Supplemental Figure 1. ALS patient demographics and antibody confirmation. (A, B and C) We initially stratified the human ALS spinal cord patient tissues used in this study based upon diagnosis (sporadic vs. familial), age and sex. (D and E) Before probing patient tissue, we first confirmed that the N protein rabbit polyclonal antibody recognized N protein using commercially available slides of SARS-CoV-2 infected lung tissue using either routine DAB immunohistochemistry (IHC) (brown staining) or confocal imaging IF (red fluorescence). Purple arrows indicate N protein reactivity in lung cells. Scale bars = 20.0μm. (F and G) We also assessed control and patient tissues for ALS pathology by probing with TDP-43 and assessing its localization and cytoplasmic inclusion status using IHC (Brown staining). Black arrows indicate TDP-43 reactivity. Note the nuclear staining profile in control tissue, and the presence of large cytoplasmic, inclusions in ALS tissue. Scale bar = 20μm.

Supplemental Figure 2. Schematics of the optogenetic system used in this study. Cytoplasmic, diffuse Opto-NCP self-associates into BMCs with BL stimulation. This process is reversible upon the removal of BL. Cry2Clust optogene tag (Blue), N protein (Purple) and mCherry fluorescent tag (Red). (B) Schematic of the optogenetic methodology used in cell culture for both fixed and live cell imaging. Both schematics were created in BioRender.

Supplemental Figure 3. mCh-Opto and NCP-aCh do not colocalize with TDP-43. (A) mCh-Opto robustly forms coarse granular cytoplasmic condensates with Blue light (BL) stimulation (10,000 lux, 465 nm), as well as with ambient light contamination (No stimulation; NS). Thus, ambient light must be controlled for in experimentation. In both paradigms, mCh-Opto condensates do not colocalize with TDP-43. Scale bars = 10.0µm (B) Merged images underwent 3D deconvolution to remove background and improve the visual specificity of colocalization. Scale bars = 5.0μm. (C) Transient expression of NCP-aCh overnight (16 hours) does not form cytoplasmic BMCs, nor does it colocalize with TDP-43.

Supplemental Figure 4. I*n vitro* Opto-NCP Colocalizes with ALS-Associated Cytoplasmic RNA Binding Proteins. Blue light induced Opto-NCP BMCs colocalize with PABPC1 (PCC = 0.47) (A and B), G3BP1 (PCC = 0.43) (C and D), and TIA1 (PCC = 0.43) (E and F). BL = Blue light stimulation. NS = No stimulation. Yellow box = formation of N protein BMCs and colocalization. Scale bars = 10.0μm. Merged images underwent 3D deconvolution to remove background and improve the visual specificity of colocalization. Scale bars = 5.0μm, 2.0μm and 0.5μm.

Supplemental Figure 5. BL Stimulation does not cause cell stress. Neither eIF2S1 (A) or phospho-eIF2S1 (B) colocalized with G3BP1-immunoreactive Opto-NCP condensates. BL = Blue light stimulation. NS = No stimulation. Yellow box = formation of NCP BMCs and signal overlap with G3BP1. Scale bars = 10.0µm (C) A Comet assay, measuring DNA damage in individual cells, shows minimal DNA tail formations in non-stimulated and BL stimulated samples. Samples were either non-transfected or transiently transfected with NCP-aCh or Opto-NCP. Samples were exposed to six hours of BL stimulation to chronically saturate samples with BL. White dashed lines outline the nucleus and the Green fluorescent DNA tails. Scale bars = 25µm. (D and E) Measurement of DNA tail length and tail DNA percentage show non-significance between non-stimulated and BL stimulated samples.

