## Supplemental Tables for "SARS-CoV-2 Nucleocapsid Protein forms biomolecular condensates and interacts with ALS-associated RNA binding proteins *in vitro*"

Supplemental Table 1. ALS Patient Tissue Demographics and Disease Features

| Case | Sex | Age at Death (Years) | Duration (Months) | Site of Onset | ALS-Specific Genotyping | Neuropathological Diagnosis | N-protein Positive |
|---|---|---|---|---|---|---|---|
| 1 | M | 56 | 22 | Bulbar | C9*orf*72** | fALS - TDP | + |
| 2 | M | 66 | 23 | Unknown | C9*orf*72 | fALS- TDP | - |
| 3 | M | 63 | 24 | Dementia / Bulbar | C9*orf*72 | fALS FTLD | + |
| 4 | F | 54 | 6 | Limb | SOD1 A4T | fALS | + |
| 5 | F | 48 | 60 | Limb | SOD1 I113T | fALS | - |
| 6 | F | 85 | 96 | Bulbar | Negative | FTLD (PSP) - Tau | + |
| 7 | M | 62 | 24 | Respiratory | Negative | sALS | - |
| 8 | F | 76 | 48 | Dementia | C9*orf*72 | fALS FTLD - TDP | - |
| 9 | F | 66 | 204 | Limb | SOD1 D90A | fALS | + |
| 10 | M | 54 | 15 | Limb | C9*orf*72 | fALS | - |
| 11 | F | 64 | 42 | Limb | C9*orf*72 | fALS AD (ABC = A1, B1, C1) | + |
| 12 | M | 66 | Unknown | Unknown | C9*orf*72 | fALS | - |
| 13 | F | 76 | 44 | Bulbar | Negative | ALS AD (ABC = A2, B2, C2) - Congophilic Angiopathy | - |
| 14 | M | 75 | 48 | Behavioural / | Negative | ALS FTLD BI - Tau | - |

| Case | Sex | Age at Death (Years) | Duration (Months) | Site of Onset | ALS-Specific Genotyping | Neuropathological Diagnosis | N-protein Positive |
|---|---|---|---|---|---|---|---|
| | | | | Bulbar | | | |
| 15 | M | 60 | 29 | Bulbar | Negative | sALS | - |
| 16 | F | 66 | 66 | Bulbar | Negative | sALS | - |
| 17 | F | 46 | 48 | Limb | Negative | sALS | - |
| 18 | F | 56 | 24 | Limb | Negative | sALS | + |
| 19* | M | 55 | 46 | N/A | Negative | Pick's Disease | - |
| 20 | F | 66 | 21 | Limb | C9*orf*72 | ALS (PART) | - |
| 21 | F | 78 | 8 (MAID) | Bulbar | Negative | ALS alpha synucleinopathy | - |
| 22 | F | 65 | 72 | Bulbar | Negative | sALS | - |
| 23 | M | 79 | 126 | Leg | Unknown/Not Available | sALS | + |
| 24 | M | 76 | 64 | Right Arm | Unknown/Not Available | sALS | + |
| 25 | F | 76 | 216 | Speech | Unknown/Not Available | sALS | - |
| 26 | M | 84 | 48 | Left Leg | Unknown/Not Available | sALS | - |

| Case | Sex | Age at Death (Years) | Duration (Months) | Site of Onset | ALS-Specific Genotyping | Neuropathological Diagnosis | N-protein Positive |
|---|---|---|---|---|---|---|---|
| 27 | M | 59 | 16 | Right Arm | Unknown/Not Available | sALS | - |
| 28 | F | 71 | 24 | Right Leg | Unknown/Not Available | sALS | - |
| 29 | M | 72 | 63 | Bilateral Arms | Unknown/Not Available | sALS | - |
| 30 | M | 70 | 20 | Bulbar | Unknown/Not Available | sALS | - |
| 31* | M | 68 | -- | -- | Negative | Sarcoma | - |
| 32* | M | 67 | -- | -- | Negative | Cerebral Vascular Disease | - |
| 33* | M | 61 | -- | -- | Negative | Transplant | - |
| 34* | F | 53 | -- | -- | Negative | Heart disease | - |
| 35* | F | 53 | -- | -- | Negative | Liver failure | - |

* Denotes control cases
**Pathogenic hexanucleotide expansion

Abbreviations:
AD: Alzheimer's disease
ALS: Amyotrophic Lateral Sclerosis
fALS: Familial ALS
sALS: Sporadic ALS
BI: Behavioral impairment
FTLD: Frontotemporal Lobar Degeneration
MAID: Medical assistance in dying
OPTN: Optineurin
PART: Primary age-related tauopathy
PSP: Progressive Supranuclear Palsy
SOD1: Superoxide Dismutase 1
Tau: Microtubule-Associated Protein Tau
TDP: TAR DNA-binding protein 43
VUS: Variant of unknown significance

Supplemental Table 2. Primer Information

| Target | Application | Primer | Sequence |
|---|---|---|---|
| *TARDBP* | RT-PCR & qPCR | Fwd<br>Rev | TCAGGGCCTTTGCCTTTGTT<br>TGCTTAGGTTCGGCATTGGAT |
| *POLDIP3* | RT-PCR & qPCR | Fwd<br>Rev | GCTTAATGCCAGACCGGGAGTTG<br>TCATCTTCATCCAGGTCATATAAATT |
| *GAPDH* | RT-PCR & qPCR | Fwd<br>Rev | TCGCCAGCCGAGCCA<br>GAGTTAAAAGCAGCCCTGGTG |
| NCP-aCh | DNA Assembly Cloning | Fwd<br>Rev | TTAGTGAACCGTCAGATCCGCTAGCATGTCCGACAACGGTCCCCAGAACCAG<br>ATGGCTGATTATGATCTAGAGTCGCTTACTTGTACAGCTCGTCCATGCCG |
| Opto-NCP | DNA Assembly Cloning | Fwd 1<br>Rev 1<br>Fwd 2<br>Rev 2<br>Fwd 3<br>Rev 3 | TTAGTGAACCGTCAGATCCGCTAGCATGGTGAGCAAGGGCGAG<br>GGGACCGTTGTCGGATGCTCCAGAAGATCCCTTGTACAGCTCGTCCATGC<br>CATGGACGAGCTGTACAAGGGATCTTCTGGAGCATCCGACAACGGTCCC<br>CTTTTTGTCCATCTTTGCACCACTACTACCAGCCTGGGTGGAGTCAGC<br>GCTGACTCCACCCAGGCTGGTAGTAGTGGTGCAAAGATGGACAAAAAG<br>TGATTATGATCTAGAGTCGCTTAGTTATCTAGATCCGGTG |
| Opto-$NCP_{M210}$ | DNA Assembly Cloning | Fwd 1<br>Rev 1<br>Fwd 2<br>Rev 2 | GAACCGTCAGATCCGCTAGCATGGTGAGCAAGGGCGAG<br>GTCACCTCCGTTGCCGGCCGCTCCAGAAGATCCCTTGTACAG<br>TGTACAAGGGATCTTCTGGAGCGGCCGGCAACGGAGGTGA<br>TGATTATGATCTAGAGTCGCTTAGTTATCTAGATCCGGTGGATCCCG |

| Target | Application | Primer | Sequence |
|---|---|---|---|
| Opto-NCP$_{M234}$ | DNA Assembly Cloning | Fwd 1 | GAACCGTCAGATCCGCTAGCATGGTGAGCAAGGGCGAG |
| | | Rev 1 | TGCTGACCCTTTCCGGACGCTCCAGAAGATCCCTTGTACAG |
| | | Fwd 2 | TGTACAAGGGATCTTCTGGAGCGTCCGGAAAGGGTCAGCAG |
| | | Rev 2 | TGATTATGATCTAGAGTCGCTTAGTTATCTAGATCCGGTGGATCCCG |
| Opto-NCP$_{N\text{-}Term}$ | DNA Assembly Cloning | Fwd 1 | CCGCCACTCCACCGGCGGCATGGACGAGCTGTACAAGGGATCTTCTGGAGCATCCGACAACGGTCC |
| | | Rev 1 | TGCACCACTACTACCACGAGCTGGGCTGGTGCCTCTGGAGCTTCC |
| | | Fwd 2 | ACCAGCCCAGCTCGTGGTAGTAGTGGTGCAAAGATGGACAAAAAGAC |
| | | Rev 2 | TCTCTCATGCCAGCATCCACGAGTGGGTACCCTGTCCTTCCCTGTCTCCATGCCTTGAAT |
| pUltra-Opto-NCP | DNA Assembly Cloning | Fwd | TGGGCTGCAGGTCCGATCCAGCTAGCATGGTGAGCAAGGGCGAGGAGGAT |
| | | Rev | GAGTCGACGACTCCGGAACGTTAGTTATCTAGATCCGGTGGATCCCGGGC |
