## Supplemental Figures for "SARS-CoV-2 Nucleocapsid Protein forms biomolecular condensates and interacts with ALS-associated RNA binding proteins *in vitro*"

Supplemental Figure 1.

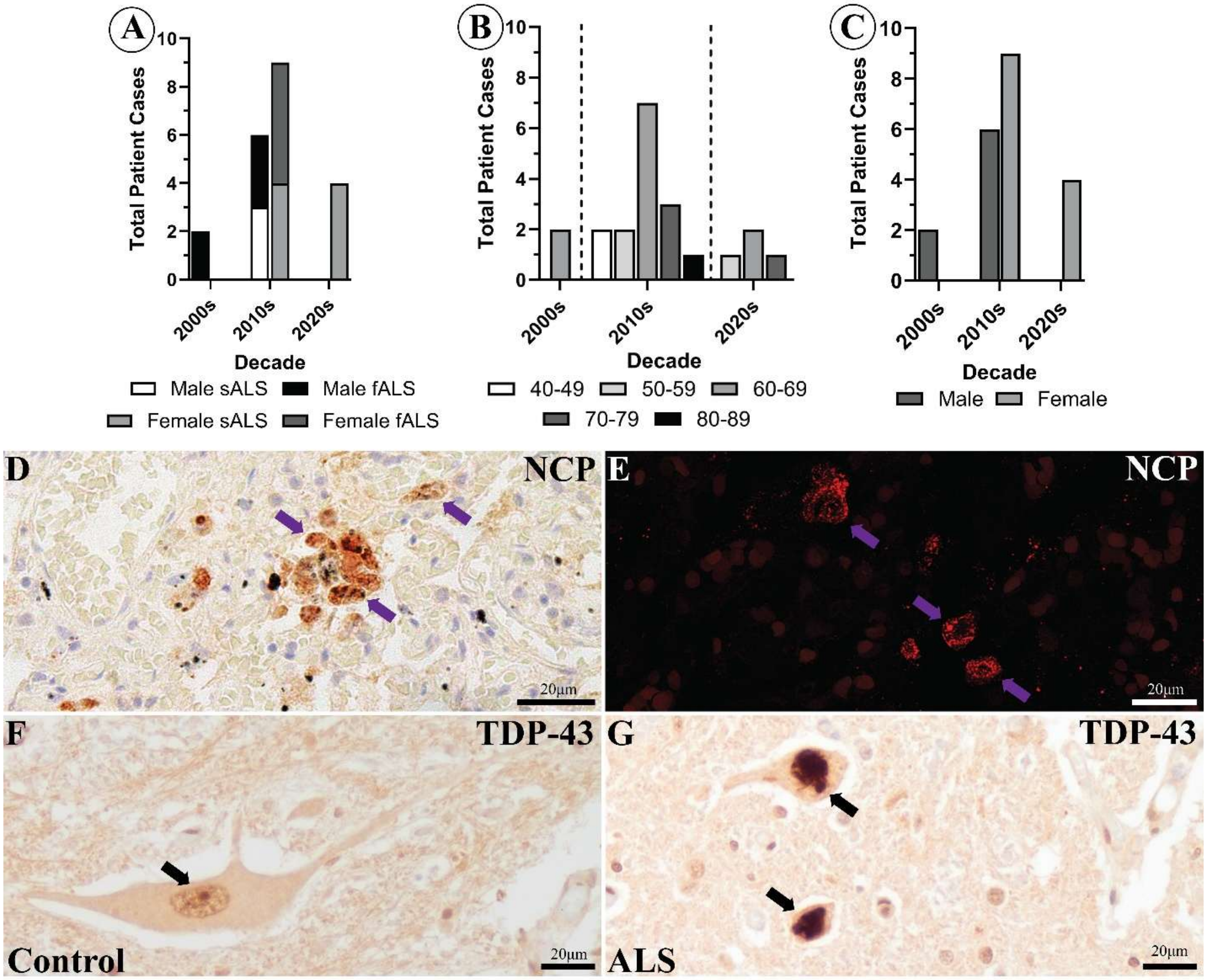

Supplemental Figure 2.

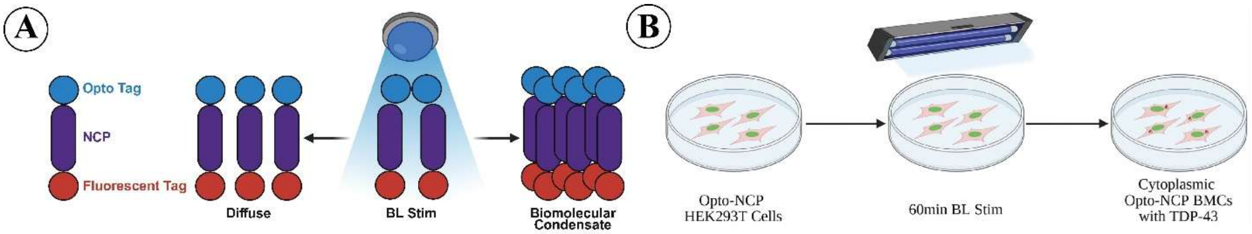

Supplemental Figure 3.

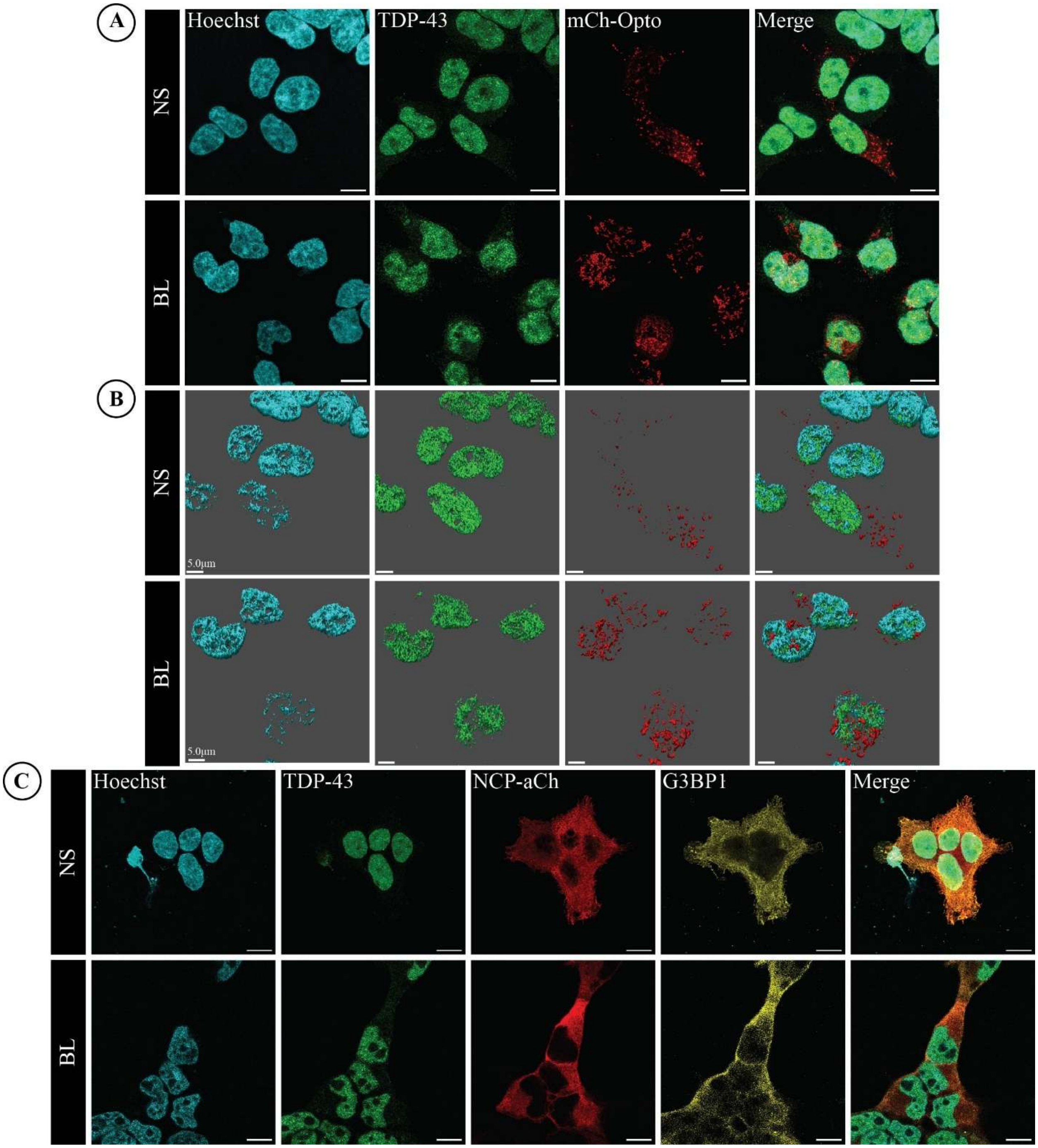

Supplemental Figure 4

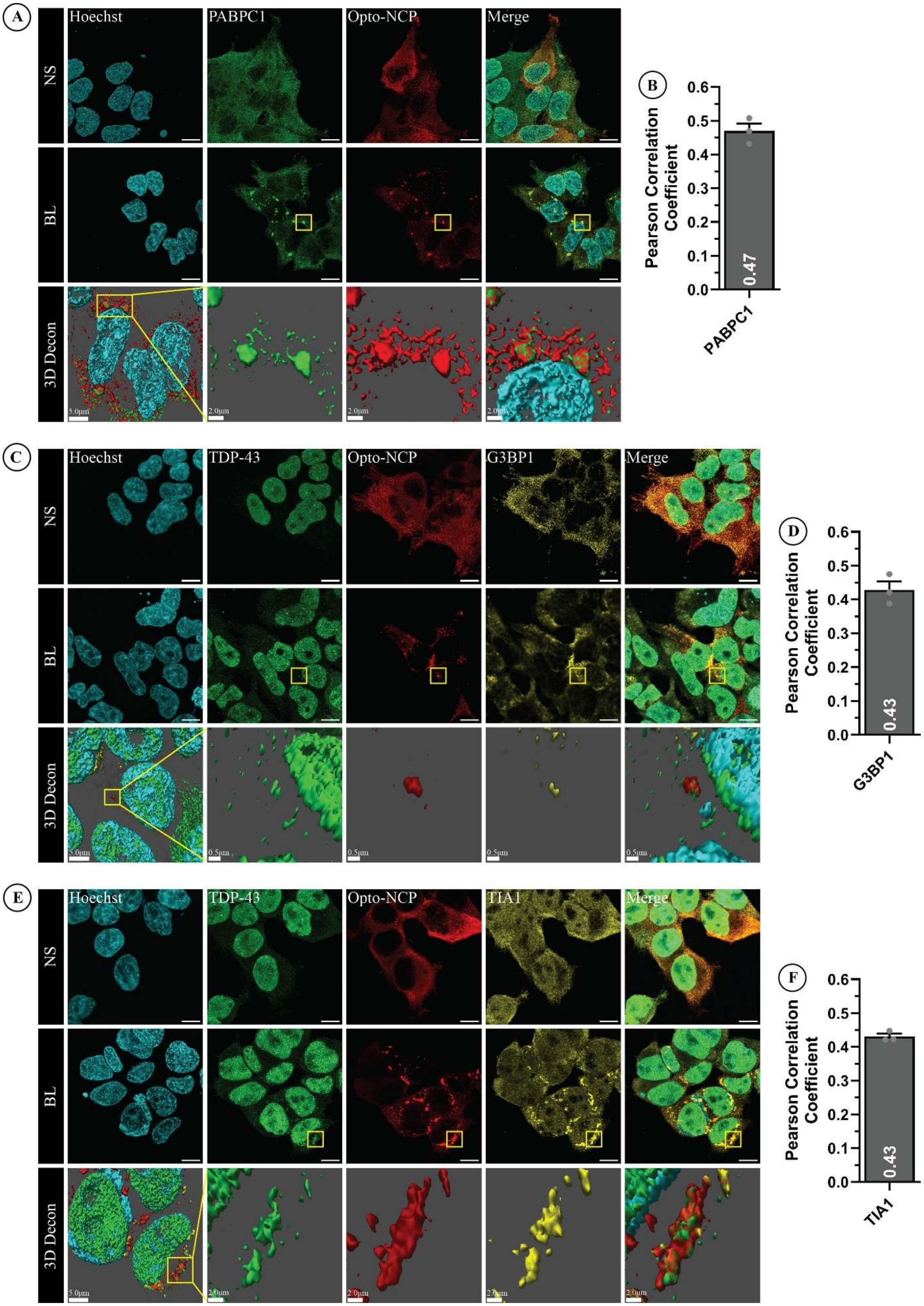

Supplemental Figure 5.

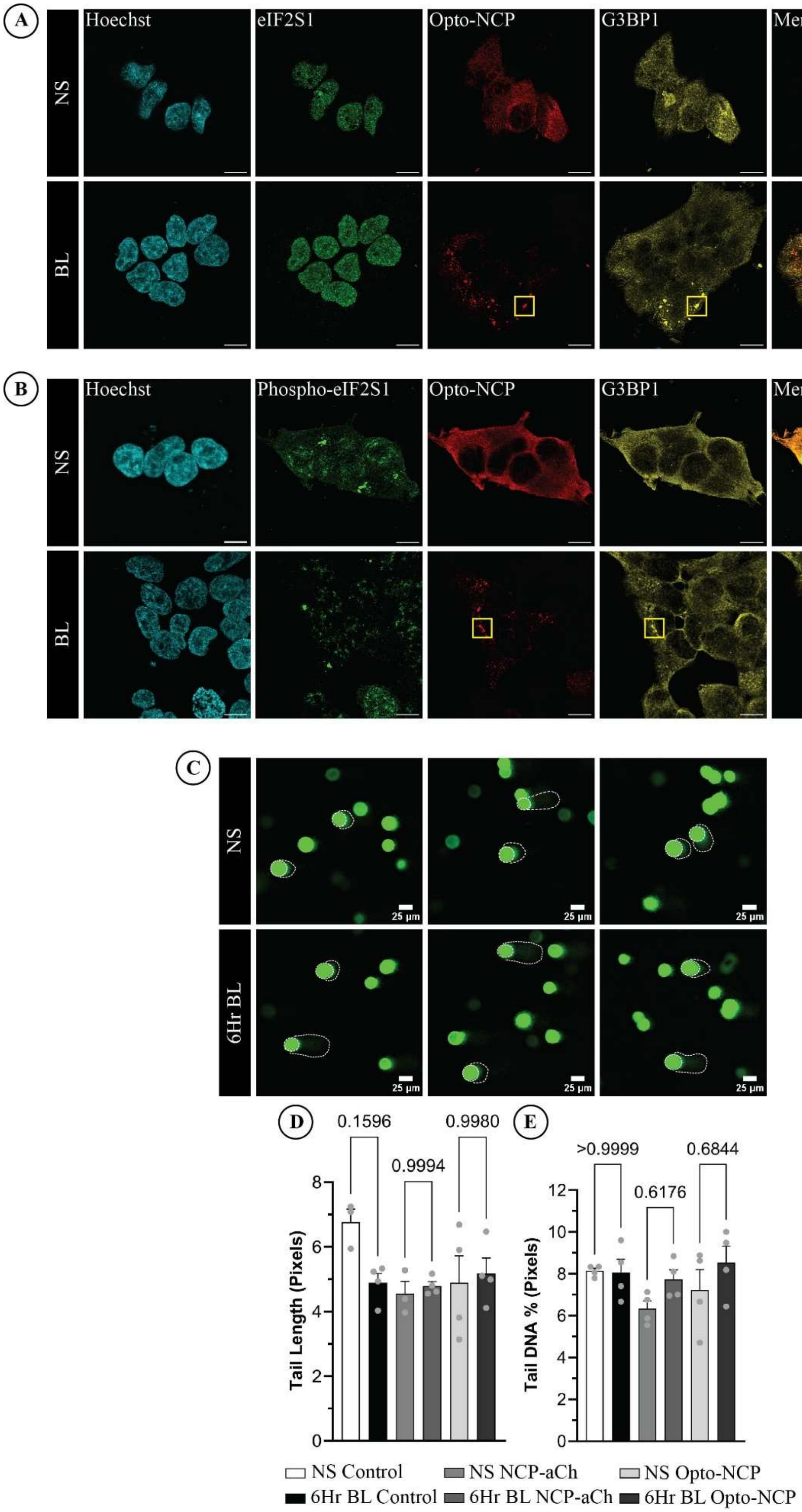
